# Microglial amyloid beta clearance is driven by PIEZO1 channels

**DOI:** 10.1101/2022.03.18.484831

**Authors:** Henna Konttinen, Valeria Sitnikova, Yevheniia Ishchenko, Anastasia Shakirzyanova, Luca Giudice, Irene F Ugidos, Mireia Gómez-Budia, Nea Korvenlaita, Sohvi Ohtonen, Irina Belaya, Feroze Fazaludeen, Nikita Mikhailov, Maria Gotkiewicz, Kirsi Ketola, Šárka Lehtonen, Jari Koistinaho, Katja M Kanninen, Damian Hernández, Alice Pébay, Rosalba Giugno, Paula Korhonen, Rashid Giniatullin, Tarja Malm

## Abstract

**Background:** Microglia are the endogenous immune cells of the brain and act as sensors of pathology to maintain brain homeostasis and eliminate potential threats. In Alzheimer’s disease (AD), toxic amyloid beta (Aβ) accumulates in the brain and forms stiff plaques. In late-onset AD accounting for 95% of all cases, this is thought to be due to reduced clearance of Aβ. Human genome-wide association studies and animal models suggest that reduced clearance results from aberrant function of microglia. While the impact of neurochemical pathways on microglia have been broadly studied, mechanical receptors regulating microglial functions remain largely unexplored.

**Methods:** Here we showed that a mechanotransduction ion channel, PIEZO1, is expressed and functional in human and mouse microglia. We used a small molecule agonist, Yoda1, to study how activation of PIEZO1 affects AD-related functions in human induced pluripotent stem cell (iPSC) -derived microglia-like cells (iMGL) under controlled laboratory experiments. Cell survival, metabolism, phagocytosis and lysosomal activity were assessed using real-time functional assays. To evaluate the effect of activation of PIEZO1 *in vivo*, 5-month-old 5xFAD male mice were infused daily with Yoda1 for two weeks through intracranial cannulas. Microglial Iba1 expression and Aβ pathology were quantified with immunohistochemistry and confocal microscopy. Published human and mouse AD datasets were used for in-depth analysis of *PIEZO1* gene expression and related pathways in microglial subpopulations.

**Results:** We show that PIEZO1 orchestrates Aβ clearance by enhancing microglial survival, phagocytosis, and lysosomal activity. Aβ inhibited PIEZO1-mediated calcium transients, whereas activation of PIEZO1 with a selective agonist, Yoda1, improved microglial phagocytosis resulting in Aβ clearance both in human and mouse models of AD. Moreover, PIEZO1 expression was associated with a unique microglial transcriptional phenotype in AD as indicated by assessment of cellular metabolism, and human and mouse single cell datasets.

**Conclusion:** These results indicate that the compromised function of microglia in AD could be improved by controlled activation of PIEZO1 channels resulting in alleviated Aβ burden. Pharmacological regulation of these mechanoreceptors in microglia could represent a novel therapeutic paradigm for AD.

**GRAPHICAL ABSTRACT:** 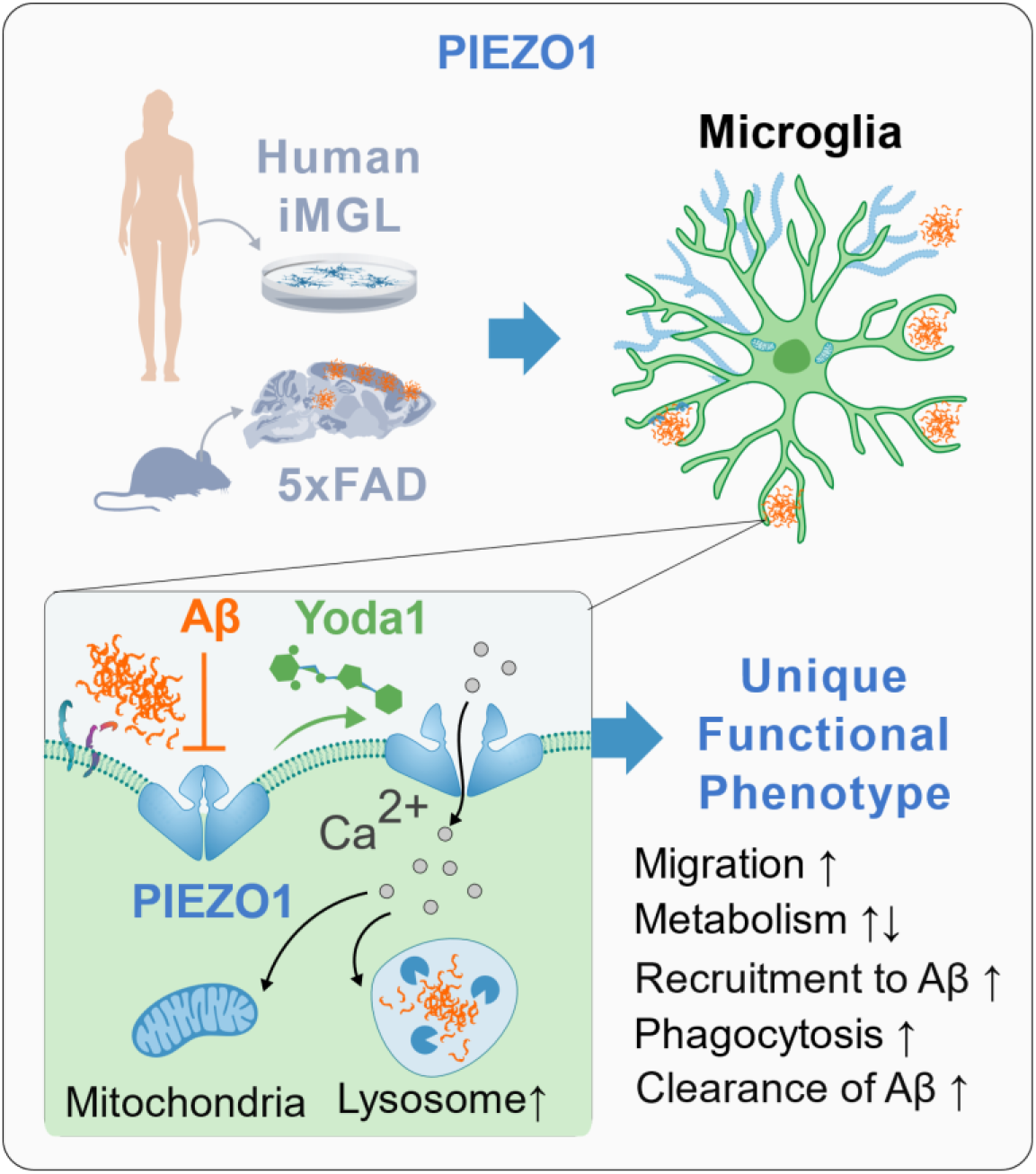

## BACKGROUND

Microglia are dynamic immune cells that chemically and mechanically interact with their environment in the brain. They continuously survey the brain parenchyma, provide trophic support for neurons, remove unnecessary synapses and clear foreign materials through phagocytic and cytotoxic mechanisms(1,2). Microglia act as antigen-presenting cells and contribute to brain homeostasis by secreting a plethora of pro- and/or anti-inflammatory cytokines and other signaling molecules depending on the situational cues(1,2). Genome-wide association studies (GWAS) implicate dysfunction in several microglial innate immunity genes to increase the risk for Alzheimer’s disease (AD). In addition, late-onset AD (LOAD), covering the majority of AD cases, is associated with impairment in the clearance of amyloid beta (Aβ)(3,4), whereas rare genetic early-onset AD (EOAD) is more evidently caused by increased production of Aβ by neurons. The inability of microglia to clear accumulating toxic Aβ together with their proinflammatory functions are thought to be a significant contributor to LOAD pathology(5,6). Thus, reshaping microglial functions represents a promising strategy for clearing accumulating deposits in AD and other neurodegenerative diseases with aberrant aggregates.

With the progression of AD pathology, aggregating Aβ form local stiff plaque deposits, thereby creating a stark contrast for the soft brain tissue (Aβ∼3 × 109 Pa vs normal brain ∼200–500 Pa)(7–10). *In vitro*, microglia are attracted to stiff regions, and they upregulate inflammatory mediators and change their morphology on stiffer substrates(11,12). *In vivo*, implantation of stiff foreign bodies enhances microglial activation, ultimately leading into the encapsulation of the foreign body(12), resembling the manner in which microglia envelope the amyloid plaques(13). This adaptation to mechanical stimuli suggests the presence of specific mechanosensors in microglia. The recently described PIEZO ion channels are amongst the most specific and sensitive mechanotransducers that translate extracellular mechanical forces to intracellular molecular signaling cascades(14,15). PIEZO2 channels are mainly expressed in the nociceptive system(16–18), while PIEZO1 channels are expressed in neurons and in non-neuronal cell types in various regions of the brain(19,20). It has been demonstrated that soluble Aβ prevents PIEZO1-mediated Ca^2+^ influx in HEK293 cells(21), and that astroglia upregulate PIEZO1 around extracellular Aβ plaques(19), suggesting a link between AD pathology and the function of brain cells that express these channels. However, the functional role of PIEZO channels in microglia remains unexplored.

Here we report that PIEZO1 is expressed in mouse microglia and human induced pluripotent stem cell (iPSC)-derived microglia-like cells (iMGLs). The activation of PIEZO1 with Yoda1, a small molecule agonist of the PIEZO1 channel, induces Aβ clearing functions in human iMGLs. Moreover, treatment of the 5xFAD mouse model of AD with Yoda1 recruits microglia towards Aβ plagues and leads to Aβ clearance in the hippocampus and cortex. In agreement with a previous study in HEK293 cells (21), we confirm that Aβ inhibits PIEZO1 and demonstrate this for the first time in human iMGLs. Supporting our findings, existing transcriptional datasets indicate that the expression of *PIEZO1* is altered in specific disease-related subpopulations of microglia in the human AD and 5xFAD brains. The main findings are illustrated in a graphical abstract.

## RESULTS

### 1. Human Microglia express highly mechanosensitive PIEZO1 channels

We characterized the expression of PIEZO channels in microglia across species in human iMGLs, mouse primary microglia and secondary microglial cell lines, as well as in transcriptional databases. Murine primary microglia and the BV2 cell line expressed *Piezo1* abundantly compared to primary rodent trigeminal neurons(50) and astrocytes(51) that were used as positive controls (Fig. 1A). To study human microglia, we differentiated human iPSC lines (Table 1) into iMGLs that resemble human microglia in their morphology, marker expression, functional aspects and capacity to respond to ATP and ADP with Ca^2+^ transients(52). Human iMGLs and human microglial SV40 cell line expressed *PIEZO1* (Fig. 1A) whereas *PIEZO2* was only detected in the SV40 cell line and primary mouse microglia (Fig.S1A). We validated this by using published human(53) and mouse(54) RNA-seq datasets (GSE52564(54), GSE73721(53), phs001373.v1.p1(55), GSE125050(56), syn18485175(57), GSE99074(58); Fig. S1B-D). To verify further the expression of *PIEZO1/2* in iMGLs, we compared two published RNA-seq datasets of iMGLs produced similarly (GSE135707(52))* or analogous(GSE133433(37)) to our method. In both datasets, iMGLs expressed *PIEZO1* more abundantly than *PIEZO2* (Fig. 1B). The levels of the two genes in iMGLs resembled the expression in fetal and adult human microglia, whereas iPSCs expressed *PIEZO2* at a higher level and induced hematopoietic progenitor cells (iHPCs), CD14 and CD16 positive monocytes (CD14M, CD16M), and dendritic cells (DC) had higher *PIEZO1* expression (Fig. 1B). Immunocytochemistry corroborated prominent localization of PIEZO1 channels on the cellular membrane both in human iMGLs (Fig. 1C, Fig. S1E) and mouse microglia (Fig. 1D).

**Table 1.** Data for human iPSC cell lines. All with APOE3/3 alleles and derived from fibroblasts of skin biopsies.

| iPSC line | Sex | Health status | Age years | Method | Karyo-type | Ref | Assays |
| --- | --- | --- | --- | --- | --- | --- | --- |
| **MAD1 clone 7** | M | Healthy | 67 | Sendai Virus 2.0 | 46 XY Normal | Manuscript under prep. | Ca-imaging, LDH, MTT, Mitostress, Migration, pHrodo/Aβ phagocytosis |
| **MAD6 clone 1** | M | Healthy | 63 | Sendai Virus 2.0 | 46XY Normal | Fagerlund et al. 2022 | Ca-imaging, LDH, MTT, Mitostress, LysoView, Migration, pHrodo/Aβ phagocytosis |
| **MAD8 clone 1** | M | Healthy | 64 | Sendai Virus 2.0 | 46 XY Normal | Manuscript under prep. | Ca-imaging, Cytotox, LDH, MTT, Mitostress, pHrodo/Aβ phagocytosis |
| **Ctrl 8.2** | F | Healthy | Adult | Sendai Virus 1.0 |  | Holmqvist et al. 2016 | Cytotox |
| **MBE2968 clone 1** | F | Healthy | 65 | Episomal nucleofection | 46XX Normal | Konttinen et al. 2019 | All assays, Ca-imaging, Cytotox, LDH, MTT, Mitostress, CBA, LysoView, pHrodo/Aβ phagocytosis |
| **MBE2960 polyclonal** | M | Healthy | 78 | Episomal nucleofection | 46XY Normal | Munoz et al. 2020 | Ca-imaging, LysoView, Mitostress |
| **TOB0644_F_B9_F3** | M | Healthy | 73 | Episomal nucleofection, CRISPR edited from *APOE4/4* | 46XX Normal | Hernandez et al. 2021 | Ca-imaging |
| **LL190 clone 1.5** | F | Healthy | 44 | Sendai Virus 2.0 | 46XX Normal | Oksanen et al. 2017 | ICC |
| **Isogenic AD4 clone 1.6.12.10** | F | Corrected presymptomatic *PSEN1DE9* (EOAD) | 47 | Sendai Virus 2.0, CRISPR corrected *PSEN1DE9* | 46XX Normal | Oksanen et al. 2017 | Ca-imaging, CBA |
| **Isogenic AD4 clone 1.5.6.1** | M | Corrected *PSEN1DE9* (EOAD) | 48 | Sendai Virus 2.0, CRISPR corrected *PSEN1DE9* | 46XY Normal | Oksanen et al. 2017 | Ca-imaging, CBA |
| **BIONI010-C-2** | M | Healthy | 15-19 | Non-integrating episomal, CRISPR edited from *APOE4/4* | 46 XY Normal | EBISC | ICC, LysoView, pHrodo |

CBA, cytokine bead array; EOAD, early-onset Alzheimer's disease; ICC, immunocytochemistry; LDH, lactate dehydrogenase; 3-(4,5-dimethylthiazol-2-yl)-2,5-diphenyltetrazolium bromide, metabolic assay; *PSEN1DE9*, Presenilin 1 delta exon 9 deletion.

**Table 2.** List of TaqMan primer assay mixes used for mRNA expression analysis.

| Target | Gene | TaqMan Assay ID |
| --- | --- | --- |
| **Endogenous control** | *18s* | Hs99999901_s1 |
|  | *ACTB* | Hs99999903_m1 |
|  | *GAPDH* | Hs02758991_g1 |
|  |  | Hs99999905_m1 |
| **Target genes** | *PIEZO1* | Hs00207230_m1 |
|  | *PIEZO2* | Hs00401026_m1 |
|  | *Piezo1* | Mm01241549_m1 |
|  | *Piezo2* | Mm01265861_m1 |

**Table 3.** Antibodies used for immunocyto-and histochemistry.

| Antibody | Clone | Host | Dilution | Cat. No | Producer | Antigen retrieval |
| --- | --- | --- | --- | --- | --- | --- |
| **Aβ** | WO-2 | MM IgG2ak | 1:1000 | MABN10 | Sigma | 10mM sodium citrate (pH 6.0) |
| **CX3CR1** |  | RbP IgG | 1:100 | ab8021 | Abcam |  |
| **GFAP** | - | RhP IgG | 1:500 | Z0334 | Dako |  |
| **GFAP** |  | ChP IgY | 1:5000 | ab4674 | Abcam |  |
| **IBA1** | - | RbP IgG | 1:250 | 019-19741 | WAKO | 10mM sodium citrate (pH 6.0) |
| **IBA1** | GT10312 | MM IgG1 | 1:250 | MA5-27726 | Invitrogen |  |
| **P2RY12** |  | RbP IgG | 1:250 | HPA14518 | Sigma |  |
| **PIEZO1** | - | RbP IgG | 1:50 | PA5-77617 | Invitrogen |  |
| **PIEZO1** | - | RbP IgG | 1:500 | NBP1-78446 | NovusBio |  |
| **PIEZO1** | 2-10 | MM IgG | 1:200 | MA5-32876 | Invitrogen |  |
| **TMEM119** |  | RbM IgG | 1:100 | ab185333 | Abcam |  |
| **TREM2** | D8I4C | RbP IgG | 1:200 | 91068S | Cell Signaling |  |

IgG, Immunoglobulin G; IgY, immunoglobulin Y; MM, mouse monoclonal; RbM, Rabbit monoclonal; RbP, rabbit polyclonal; ChP, chicken polyclonal.

**Figure 1.**
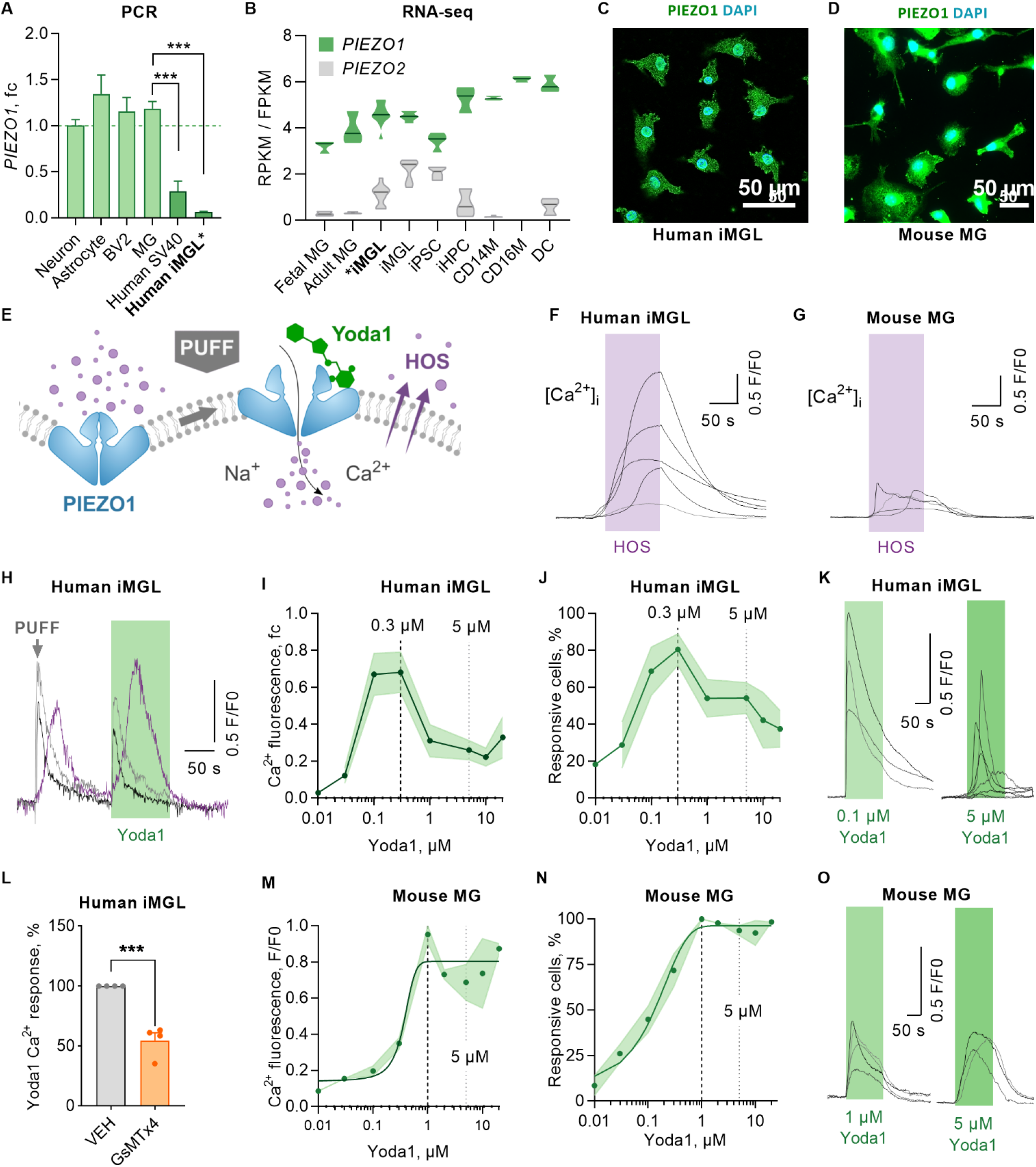
Human and mouse microglia sense mechanical forces through PIEZO1 receptor. **A** *Piezo1* gene expression in murine trigeminal neurons, astrocytes, microglia (MG) and microglial cell line (BV2); and in human microglial cell line (SV40) and iPSC-derived microglia (iMGL) analyzed by RT-qPCR (N=3-4). **B** *PIEZO1* and *PIEZO2* gene expression in human iMGLs, fetal and adult MG, iPSCs, induced hematopoietic progenitor cells (iHPCs), CD14+ and CD16+ monocytes (CD14M, CD16M), and dendritic cells (DC) obtained from human RNA-seq datasets*(21*, 28)*. N=3-6. Immunostaining of PIEZO1 (green) and nuclei DAPI (blue) in **C** iMGLs and **D** MGs. **E** Schematic for PIEZO1 activation by mechanical fluid puff, small molecule agonist Yoda1, and hypo-osmotic solution (HOS). Ca^2+^ transients of single **F** iMGLs and **G** MGs evoked by 2-minute-applications of HOS. **H** Ca^2+^ transients in iMGLs induced by mechanical fluid puffs (500 msec, 30 psi) followed with chemical activation by 0.1 µM Yoda1. n=7 coverslips. Dose-response curves for Yoda1 as **I** fold change (fc) to maximum Ca^2+^ amplitudes normalized to baseline (F/F0), **J** and as a percentage of responsive cells. Dashed line, maximum response; dotted line, 5 µM. n=2-14 with N=2-5. K Ca^2+^ transients of 0.3 µM and 5 µM Yoda1 in single iMGLs. **L** Maximum Ca^2+^ amplitudes as % compared to vehicle control (VEH) after 1 h pre-incubation with 5 µM GsMTtx4 inhibitor followed by 1-min Yoda1 application (n=4). Dose-response curves for **M** normalized maximum amplitudes and **N** percentage of responsive mouse MG (N=6). **O** Ca^2+^ transients of 1 µM and 5 µM Yoda1 in single mouse MGs. Unpaired t-test, ***p < 0.001, **p < 0.01, *p < 0.05; data repeated in n=experiments with N biological replicates. Data as mean ± SEM. See also Fig. S1 and Table 1.

Following the identification of mechanosensitive PIEZO1 channels in microglia, we next demonstrated that the iMGLs respond to two types of commonly used mechanical stimulation: 1) plasma membrane stretching caused by hypo-osmotic solution (HOS), and 2) mechanical force caused by fluid pressure puffs (Fig. 1E). Since HOS and puffs cause changes in membrane and cytoskeleton organization and thus may trigger non-specifically several mechanical receptors potentially expressed in microglia, we finally activated specifically PIEZO1 mechanotranscduction using 3) a selective small molecule agonist Yoda1 that is highly specific for PIEZO1(59). Since PIEZO1 channels are activated by conformational change upon plasma membrane stretching, we used a two-minute application of HOS to cause cell swelling. As expected, exposure to 40% lower mechano-osmotic solution with 200 mOsm/kg induced Ca^2+^ influx in iMGLs (Fig. 1F) and in primary mouse microglia (Fig. 1G). To study more fine-tuned mechanical stimulation of individual iMGLs, we created fast 500 ms pressure puffs of basic solution using a picospritzer and detected slow Ca^2+^ transients in the iMGLs (Fig. 1H). To finally verify that the Ca^2+^ transients were mediated particularly by the PIEZO1 channel, a puff was followed by an application of 0.1 µM Yoda1, a selective chemical activator of PIEZO1, demonstrating strong Ca^2+^ transients in the same cells (Fig. 1H). Dose-response curves (DRC) of Yoda1 normalized to an equal volume of vehicle (0.04% dimethylsulfoxide, DMSO) revealed high sensitivity of human microglia to Yoda1 (Fig. 1I). The maximum response was obtained in 80.5±8.5% of cells at the range of 0.1-1 µM Yoda1 while also the higher concentrations continued to induce Ca^2+^ transients, although in a smaller population of iMGLs (Fig. 1J) and with lower amplitudes (Fig. 1I, K). Conversely, inactivation of PIEZO1 suppressed Yoda1-elicited Ca^2+^ transients as illustrated by 1 h preincubation with 5 µM GsMTx-4, a mechanosensitive and stretch-activated ion channel inhibitor (Fig. 1L) (60). Mouse microglia demonstrated a more classical sigmoidal DRC and all cells were responsive to over 1 µM Yoda1 (Fig. 1M-O). In summary, these data demonstrate that across species, microglia express functional PIEZO1 channels that can be efficiently activated with the small molecule agonist Yoda1 in a species/model-dependent manner. Notably, the highest sensitivity was observed in human cells suggesting their functional role in this cell type.

### 2. Activation of PIEZO1 shapes functional phenotype of human microglia

To investigate whether PIEZO1 activation alters basic microglial functions that are important for counteracting Aβ plaques, we applied Yoda1 on iMGLs and analyzed their survival, metabolism, migration, and phagocytosis *in vitro*. First, Cytotox Green assay was used for determining a non-toxic dose of Yoda. To this end, the cells were live-imaged for 60 hours after the Yoda1 application using IncuCyte technology. As a positive control, 200 µM 1-Methyl-4-phenylpyridinium iodide was used to induce slow cell death within the experiment time window. No fresh medium was replenished to avoid mechanical activation due to media change and consequently the cells started to die after 30 hours in all wells. No differences in toxicity were observed for 0.3-20 µM Yoda1 within 24 hours as evaluated from the count of Cytotox green positive cells per confluence (Fig. 2A-B). Interestingly, 5-20 µM Yoda1 even reduced the cell death dose-dependently after 48 h compared to vehicle whereas no differences between vehicle and medium control or higher Yoda1 concentrations were observed (Fig. 2A-B, S2C). Cell confluence and cell count supported the observed cell survival induced by Yoda1 and the effect was even more visible after 60 h (Fig. 2C, Fig. S2A-B). In contrast to Piezo1 agonist, the mechanosensitive ion channel inhibitor, GsMTx4, increased cell death at 5 µM concentration (Fig. 2C-D). Since 5 µM Yoda1 was the lowest concentration to increase survival over time (Fig. 2AB, S2A-C) and to induce repeatable, although not maximal, Ca^2+^ transients (Fig. 1J-L), we used 5-20 µM doses for further functional analysis. Furthermore, the 24 h-timepoint was selected for functional studies as longer incubations would have required refreshing the culture medium. Lactate dehydrogenase (LDH) release assay confirmed Yoda1-induced protection at 5 µM concentration (Fig S2D). It was verified that 5 µM Yoda1 did not cause alterations in morphology or microglial protein expression by immunostaining for PIEZO1 and canonical microglial markers IBA1, CX3CR1, TMEM119, TREM2 and P2RY12 (Fig. S2E).

**Figure 2.**
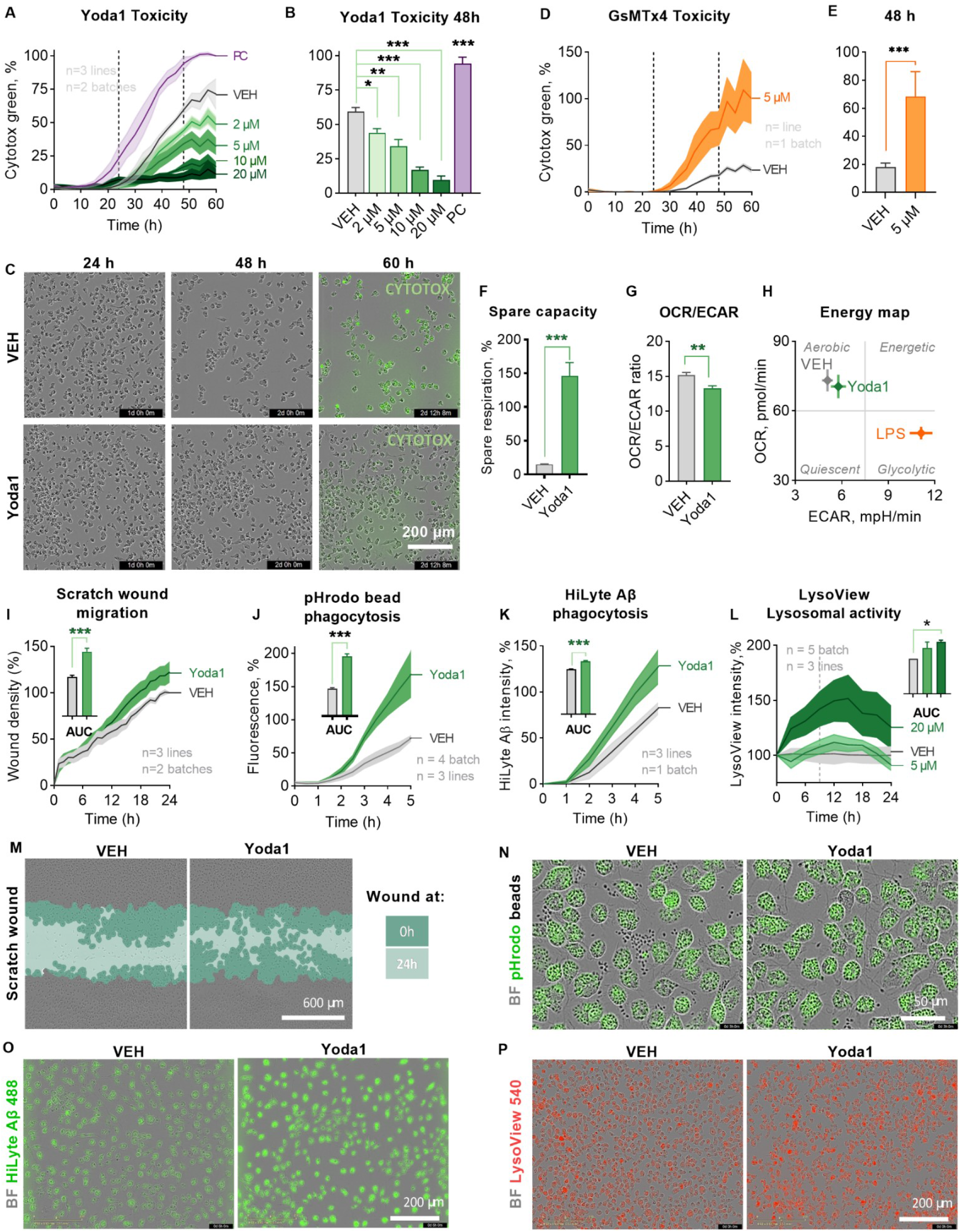
PIEZO1 orchestrates a unique immune response in human iMGLs. **A** Toxicity as count of Cytotox green cells per confluence for iMGLs treated with 2-20 µM Yoda1 and live-imaged for 60 h. Normalized to positive control (PC) of 200 µM 1-Methyl-4-phenylpyridinium iodide. N=3 in n=3. **B** Quantification at 48 h. **C** Representative images of confluency and labeling for fluorescent cytotox green reagent. **D-O** The cells were treated with 5 µM Yoda1 24 h before starting the assays. **D** Respective toxicity for iMGLs treated with 5.0 µM GsMTx4 and **E** quantification at 48 h. n=6 wells. **F** Spare respiratory capacity calculated from the oxygen consumption rate (OCR) profiles in mitostress test. n=7 with N=5. **G** Ratio of OCR to extracellular acidification (ECAR). **H** A mitostress energy map depicting OCR and ECAR compared to 20 ng/ml LPS. **I** Scratch wound density normalized to vehicle. Areas under the curves (AUC) presented as bars. N=3 in n=2. Green **f**luorescence **i**ntensity of phagocytosed **J** pHrodo beads (n=4 with N=3) and **K** 0.5 µM green HiLyte Aβ 1-42 (N=3 in n=1) per confluence over 5 h with AUC normalized to vehicle. **L** Lysosomal activity over time as intensity of pH-sensitive fluorescent red LysoView 540 reagent per confluency. n=5 with N=3. Representative images of **M** scratch wounds at 24 h overlayed with masks for the original scratches at 0 h and at 24 h, **N** cells with internalized green fluorescent pHrodo beads that show as black dots outside the cells at 3 h; **O** cells with internalized Hilyte Aβ42 488 at 5 h; and, **P** cells treated with 20 µM Yoda1 and fluorescent red LysoView 540 reagent. Two-way ANOVA with Sidak’s multiple comparisons or unpaired t-test. Significance ***p < 0.001, **p < 0.01, *p < 0.05. Data as mean ± SEM. All data repeated in n=experiments/batches with N biological replicates. **See also Fig. S2**.

We next examined whether metabolic reprogramming preceded Yoda1-induced survival as microglia change their metabolism to have adequate energetic capacity and substrates required for the stimulus-dependent protective phenotypes(52,61). The energetic profiles of iMGLs were assessed using the MitoStress Kit and a Seahorse XFe96 Analyzer (Agilent, Fig. S2F) and an increase in spare respiratory capacity was observed (Fig. 2F), as is also observed under anti-inflammatory stimulus with IL-13 and IL-4 or upon IFNγ treatment(52,61). In contrast, the ratio of OCR to ECAR indicated a shift towards more glycolytic processes (Fig. 2G), which is classically induced upon proinflammatory stimulus by lipopolysaccharide (LPS)(52,61). However, the OCR-ECAR energy map showed that Yoda1 induced only a small change in OXPHOS, whereas 20 ng/ml LPS caused a prominent shift from aerobic OXPHOS towards glycolysis (Fig. 2H, S2G-H).

Protective properties of microglia are linked to their ability to migrate to the site of pathology(62). Mechanotransduction(11,12) and overexpression of calcium-permeable PIEZO1 channels (21) have been linked to Ca^2+^-dependent processes that directly affect cell migration, including actin remodelling, myosin II contractility, binding to and integrity of the extracellular matrix. To test if PIEZO1 activation impacts the motility of iMGLs we carried out scratch wound live-imaging assays over 24 h. Quantification of the area under the curve (AUC) for the relative wound density revealed increased chemokinesis and increased wound healing after Yoda1 application (Fig. 2I and 2M). We used 100 µM adenosine diphosphate (ADP) as a positive control (Fig. 1J), since microglia increase their motility to sites of neural injury where adenosine triphosphate (ATP) is released, and ATP is rapidly converted to ADP that induces microglial migration and chemotaxis via P2Y12 receptors.

After reaching the pathological site, microglia maintain brain homeostasis by clearing damaged tissue and pathological particles through phagocytosis. Even though elevation in cytosolic Ca^2+^ is known to be required for efficient ingestion of particles and control the subsequent steps involved in the maturation of phagosome, the molecules that mediate the Ca^2+^ ion flux across the phagosomal membrane are still unknown (63). We hypothesized that PIEZO1-mediated Ca^2+^ influx would enhance phagocytosis and exposed iMGLs next to pHrodo bioparticles tagged with Zymosan ligands from yeast. The pHrodo particles are non-fluorescent at neutral pH outside the cells but fluoresce brightly in acidic pH inside phagosomes. Consistent with our hypothesis, 24-hour pre-treatment with Yoda1 increased phagocytosis quantified as green fluorescence intensity per confluency using live-cell imaging over 5 hours (Fig. 2J). This was evident in microscopy images showing fewer beads outside cells (Fig. 2N). Moreover, also the phagocytosis of AD-relevant Aβ was increased in similar experimental setting using 0.5 µM HiLyte™ Fluor 488-labeled Aβ peptides and in the quantification of green fluorescence intensity per cell area within a 5 h-follow-up period (Fig. 2K, O). After the engulfment, phagocytosis is completed in phagolysosomes that are acidic and hydrolytic organelles responsible for digesting phagocytosed cargo(64). Several AD risk genes are indicated along this pathway(64–66) and therefore we evaluated microglial lysosomal activity using red LysoView dye that accumulates in the low pH environment of the lysosomes, resulting in highly specific lysosomal fluorescence. Indeed, Yoda1 boosted lysosomal activity dose-dependently in iMGLs (Fig. 2L, P), suggesting an overall increase in the phagocytic process in microglia upon PIEZO1 activation.

### 3. Aβ inhibits the PIEZO1 channel in microglia

Ca^2+^ signaling is critical for microglial protective functions, and it is dysfunctional in AD(67–69). However, the exact mechanisms or time course of these dysfunctions have not been extensively studied. It is likely that Aβ can modulate cell membrane and cytoskeletal mechanics resulting in blockage of PIEZO1 in microglia, similarly as has been shown for HEK293 cells (21). To explore this inhibitory action in microglia, we pretreated human iMGLs with either vehicle or 1 µM soluble Aβ 1-42 for 15 and 30 minutes before applying 0.3 µM Yoda1. The most effective concentration of Yoda1 was selected based on the earlier Ca^2+^ imaging experiments (Fig.1I-J). Indeed, pretreatment with Aβ significantly dampened the Yoda1-induced Ca^2+^ influx in iMGLs (Fig. 3A-B). Since Aβ starts to fibrillize immediately after the reconstitution in water, it is likely that the soluble Aβ used in this experiment contained different forms from monomeric to fibrils, thus resembling the heterogenous forms present in the brain(70–74). It is beyond this study to identify which of the Aβ forms caused the inhibitory effect or what was the exact mechanism how Aβ inhibited PIEZO1 activity, however, the longer incubation caused more significant inhibition in Ca^2+^ transients (Fig. 3B). These results suggest that AD-related pathology may compromise microglial PIEZO1-mediated Ca^2+^ signaling and the downstream functions of microglia, providing a possible novel explanation for the failure of microglia-mediated clearance of Aβ in AD.

**Figure 3.**
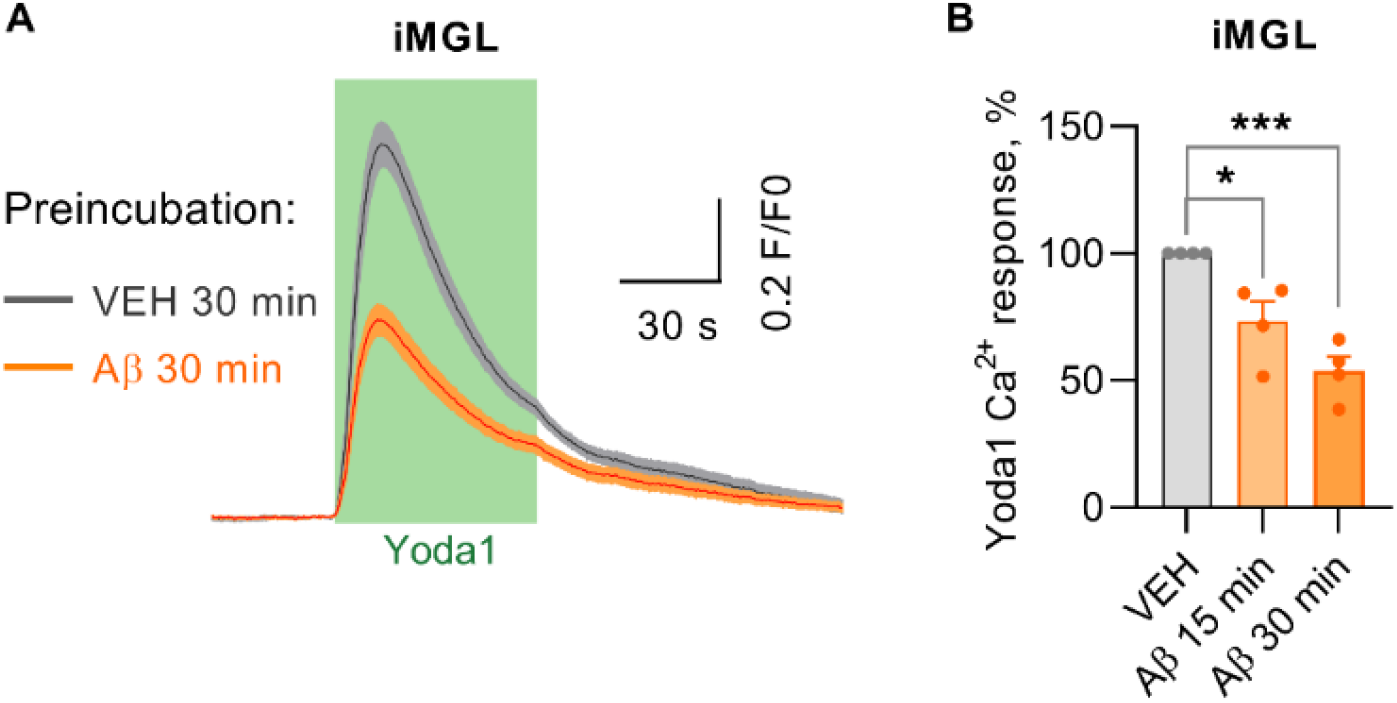
Aβ inhibits PIEZO1 activation in human iMGLs. **A** Average representative Ca^2+^ transients in iMGLs after 30 min preincubation with soluble 1 µM Aβ followed by 1-min 0.3 µM Yoda1 application from one experiment. **B** Corresponding maximum Ca^2+^ amplitudes as the % compared to VEH control. n=4 cell batches. Unpaired t-test. Significance ***p < 0.001, *p < 0.05. Data as mean ± SEM

### 4. PIEZO1 mediates clearance of Aβ *in vivo*

Since PIEZO1 activation increased microglial migration and phagocytosis, we next investigated the consequences of this potentially beneficial effect *in vivo*. First, we confirmed the presence of PIEZO1 channels in the brain of wild type (WT) and transgenic 5xFAD mice at the age of 5 months when the 5xFAD mice show prominent AD pathology with Aβ deposition and microglial activation(75,76). Immunohistochemistry illustrated abundant expression of PIEZO1 channels both in WT and 5xFAD mice and the expected upregulation of microglial IBA1 immunoreactivity in 5xFAD mice (Fig. S3A-D). Quantification of PIEZO1 immunoreactivities in hippocampi did not reveal differences between 5xFAD and WT (Fig. S3E). Similarly, no differences were observed in *Piezo1* mRNA expression using RT-qPCR (Fig. S3F).

We treated the 5xFAD mice with a daily bolus of 0.25 µg/mouse of Yoda1 or vehicle (1% DMSO5 µl bolus) 5-times per week for two weeks. The treatment was administered into the lateral ventricle using intracranially implanted cannulas for daily infusions to deliver Yoda1 in the brain (Fig 4A). To quantify the microglial distribution and the Aβ deposition we stained sagittal brain slices with IBA1 and WO2 antibodies (Fig. 4B-C). Remarkably, Yoda1-treated mice had visibly and quantifiably higher immunoreactive area of IBA1 positive microglia (Fig. 4D) and lower WO2 plaque area (Fig. 4E) in the hippocampus and cortex. To analyze the correlation of microglia and plaques in more detail, we captured 3D confocal z-stacks from hippocampi with 40x magnification (Fig. 4F-H) and revealed increased IBA1/WO2 ratio around the plaques in Yoda1 treated animals (Fig. 4I), indicating enhanced microglial clustering around the deposits. Despite these observations indicating the cleaning effect of Yoda1-activated microglia, we did not observe increased levels of PIEZO1 nor increased overlap of PIEZO1 and IBA1 (Fig S4A-E). Moreover, there was no increase in astrocytic activation as evaluated by GFAP staining or indication of altered clustering of GFAP positive cells around the WO2 plaques (Fig. S4G-M). Together these data suggest the involvement of activated microglia in the observed reduction of Aβ deposition.

**Figure. 4.**
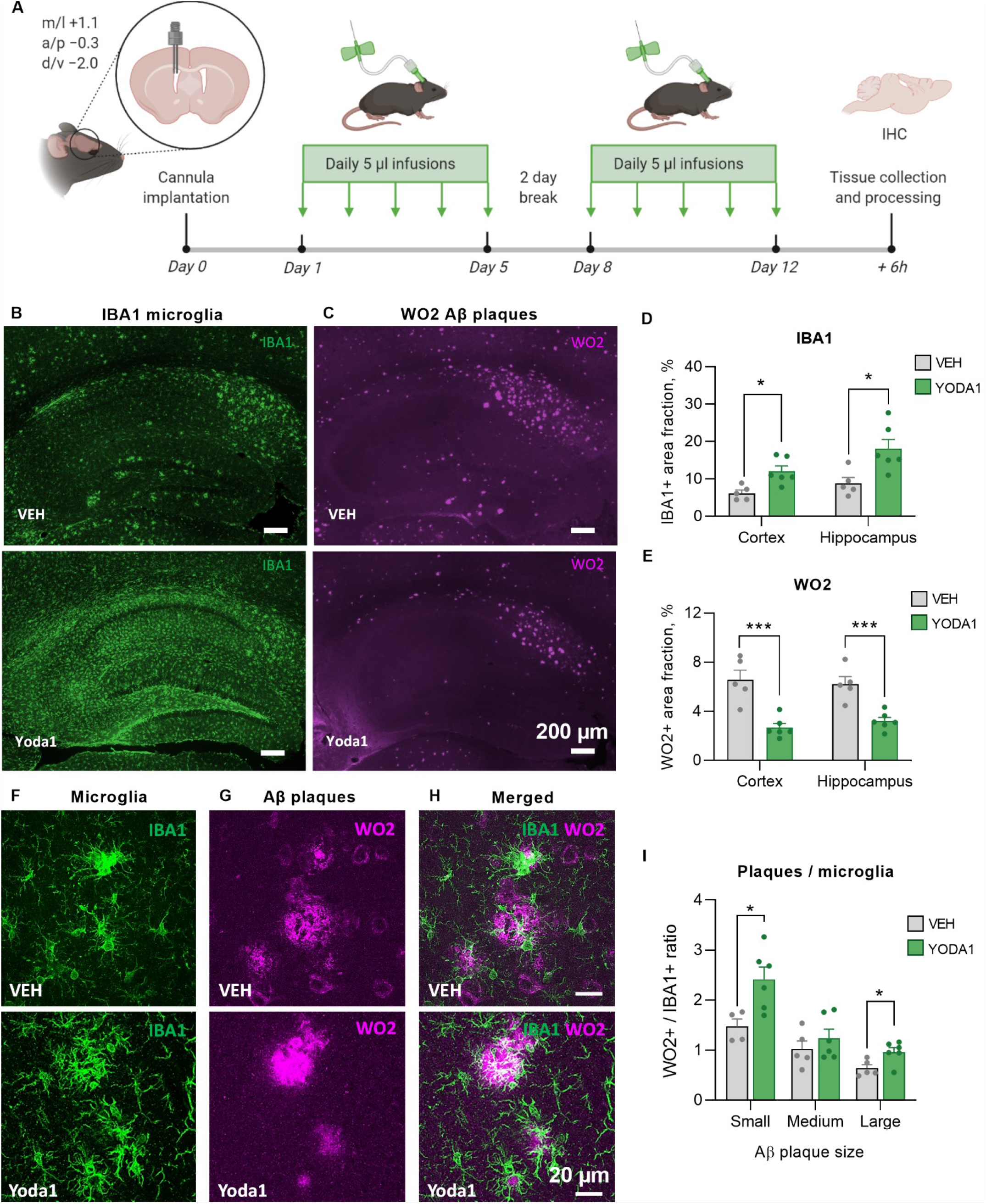
Activation of PIEZO1 mediates microglial clearance of Aβ plaques in 5xFAD mice. **A** A Schematic for *in vivo* treatments of 5-month-old 5xFAD mice with 1% DMSO in saline (VEH) or 0.25 µg Yoda1 with total of ten daily 5 µl infusions over 12-day period through intracranial ventricular cannula. Created with BioRender.com. Representative immunofluorescence images of **B** microglia (IBA1, green) and **C** Aβ plaques (WO2, magenta). Scale bar 200 µm. **D-E** Corresponding quantifications of immunoreactive area fractions in hippocampus and cortex. Representative maximum intensity projections of confocal z-stack images of double-immunostaining for **F** microglia (IBA1, green) and **G** Aβ plaques (WO2, magenta), with **H** merged images showing colocalization (white). Scale bar 20 µm. **I** Quantification of overlap of IBA1 and WO2 in small, medium, and large Aβ plaques. N=5 VEH, N=6 Yoda1 mice. Statistical outliers removed. Unpaired t-test. Significance ***p < 0.001, *p < 0.05. Data as mean ± SEM. **See also Fig. S4 and Fig. S5**.

### 5. *PIEZO1* expression is altered in AD-specific subpopulations of microglia

To better understand the role of PIEZO1 in regulating microglial subtypes in AD, we analyzed the Keren Shaul et al.(41) single-cell RNA (scRNA) and Zhou et al.(42) single nuclei (snRNA) datasets of 5xFAD mice, and Grubman *et al*. human snRNA dataset of AD patients and non-diseased age-matched individuals(40). The mouse datasets described the signature profile of disease-associated microglial (DAM) subtype in the vicinity of Aβ plaques with upregulated AD risk genes and downregulated homeostatic genes(41) and showed that the transition from homeostatic to DAM phenotype occurred through the Trem2 pathway(42). We replicated the clustering and the annotation analysis made by the authors (Fig. S5A, B, C) and depicted the *Piezo1* expressing cells in the clusters (Fig. 5A, B, C). A consensus score summarizing *Piezo1* frequency and expression in respect to the other genes revealed that microglia have the highest *Piezo1* expression, whereas in other cell types, *Piezo1* is either frequent but poorly expressed or has low frequency but is highly expressed (Fig. 5D, E, F).

**Figure 5.**
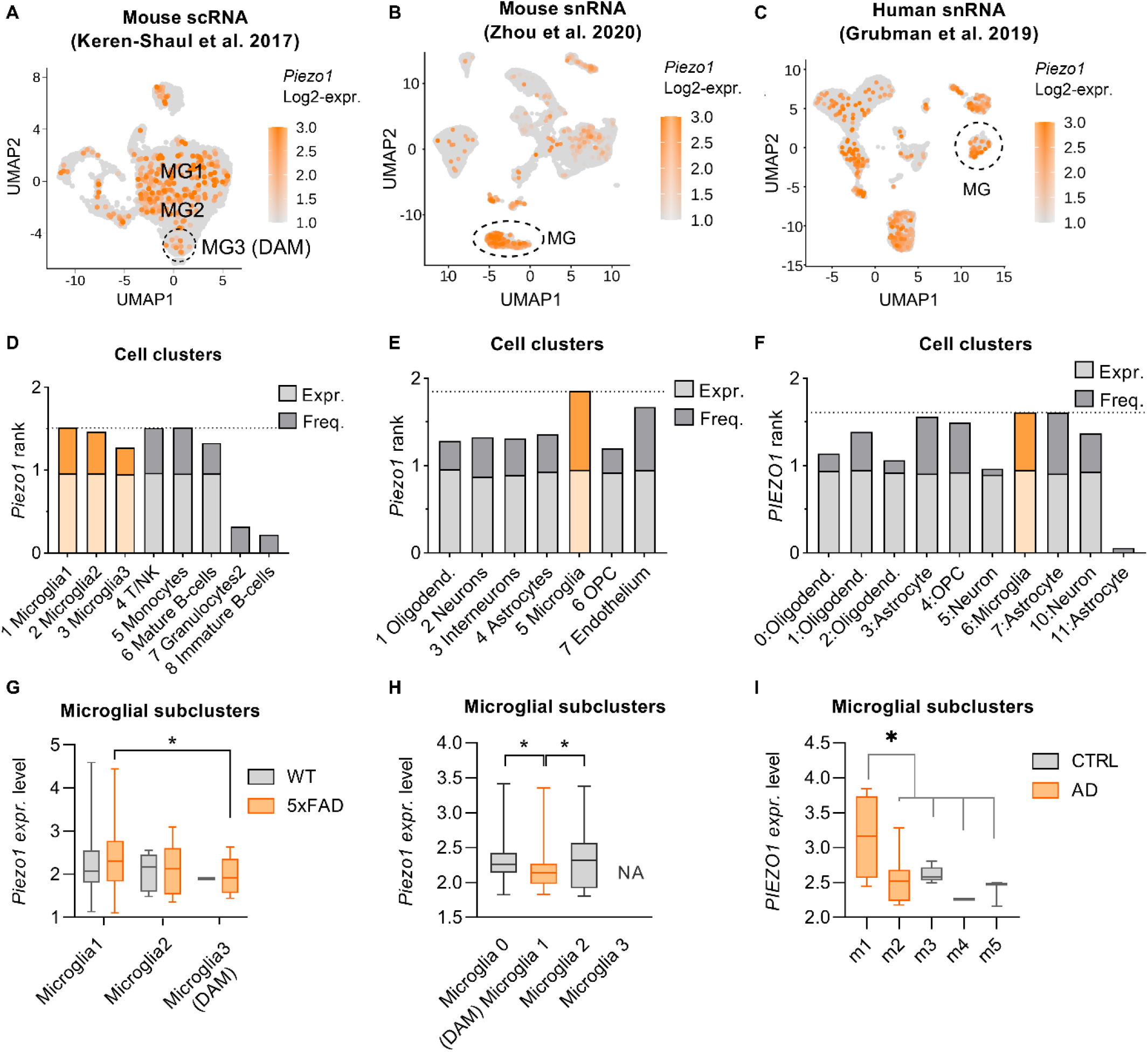
*PIEZO1* expression is altered in AD-specific subpopulations of human and murine microglia. *Piezo1* Log2 gene expression overlaid onto the Uniform Manifold Approximation and Projection (UMAP) plots of the clusters in **A** 5xFAD scRNA(41), **B** Trem2^-/-^ 5xFAD snRNA ((42)), and **C** human AD patient snRNA(40) dataset. Corresponding consensus scores summarizing the ranks of *Piezo1* frequency and expression in respect to the other genes in all annotated clusters (see Fig. S5): **D** for A data, **E** for B data and **F** for C data. Respective *Piezo1* expression level in microglial subclusters of **G** WT and 5xFAD mice in A, **H** all Trem2-/-5xFAD subclusters in B, and **I** in human microglial subtypes m1-m5 incC. Subtypes m1 and m2 are only present in AD patients *PIEZO1*. Unpaired t-test, one-way ANOVA with Dunnett’s multiple comparisons, kruskal wallis p.v. Significance *p < 0.05. Data as mean ± min and max. **See also Fig. S5**.

Supporting our hypothesis of impaired function of Piezo1 in AD, *Piezo1* was downregulated in DAM in both 5xFAD mouse datasets (Fig 5G, H) (Kruskal Wallis p.v. <= 0.05). To determine how DAM genes are regulated in Piezo1 expressing cells, we performed a correlation analysis between *Piezo1* and the DAM markers in microglial subpopulations of both datasets. Seven upregulated DAM markers (*Cd9, Ctsb, Ctsd, Ctsl, Ctsz, Fth1, Rplp2*) correlated negatively (<=-0.7) with *Piezo1*, thus while *Piezo1* decreased, they increased in DAM (Fig S5D, Table S1-2). On the contrary, thirteen downregulated DAM markers (*Cd164, Cox6a1, Glul, Malat1, Marcks, Npm1, Rpl22l1, Rpl35a, Rpl5, Rps11, Rpsa, Serinc3, Tmem173*) correlated positively, thus were also decreased in DAM similar manner as *Piezo1* (>=0.7) (Fig. S5D, Table S1-2). These genes encode proteins taking part in phagocytosis, lysosomal activity, adhesion, mitochondrial respiration, metabolism, cell shape and motility, and homeostatic microglial identity indicating the pathways related to the functions Yoda1 modulated in our microglial experiments.

Contrasting with the mouse data, human microglial subpopulations associated with AD patients (m1 and m2) had little overlap with the DAM signature(41). In agreement with this species discrepancy, we discovered that *PIEZO1* expression was upregulated in the human AD-specific subcluster m1 (Fig. 5I). Differentially expressed genes (DEGs) analysis showed, in line with the Grubman et al. 2019 data, that the m1 *PIEZO1* cells were enriched with 1) AD GWAS genes *APOC1* and *APOE*; and 2) mitochondrial genes *LINGO1, MT-ND4, MT-ND2, SNX6, MT-ATP6, MT-CYB, MT-CO2*, and *MT-CO3;* but had 3) low levels of AD GWAS gene *FRMD4A*

## DISCUSSION

Here we show a novel approach to enhance the microglial Aβ clearing function through activation of mechanotransducing PIEZO1 channels with a small molecule agonist. As we demonstrate the beneficial effects of PIEZO1 activation both in human microglia *in vitro* and in an animal model of AD *in vivo*, this calcium permeable mechanoreceptor could represent a novel translational target for LOAD patients covering the majority of AD cases associated with impairment in the clearance of Aβ (3,4) We show that functional PIEZO1 channels are expressed in human and mouse microglia and that Aβ1-42 compromises PIEZO1-mediated Ca^2+^ signaling in human iMGLs. Activation of PIEZO1 elicits a unique functional microglial state with increased phagocytosis, survival and motility with both classical pro-and anti-inflammatory metabolic features. Our in depth-analysis of single-cell sequencing datasets show that *PIEZO1* is enriched in microglial subclusters in AD patients. This correlates with the expression of certain Trem2-dependent DAM signature genes(41,42) that take part in the Aβ clearing functions in a mouse model of AD. The function of microglial cells surrounding the plaques, particularly their motility and phagocytic clearance of Aβ, may be the key determinants governing the pathological processes in AD(2,77). Importantly, it is shown here PIEZO1 activation reduced existing Aβ plaques concomitantly with the increase in microglial marker Iba1 around the plaques in 5xFAD mice *in vivo*.

Research over the last two decades has attempted to find the determinants and outcomes of the microglial behavior but their beneficial vs. detrimental contribution in AD is still not fully clear. Sustained exposure to Aβ, cytokines and other inflammatory mediators appear to cause permanent impairment in microglial function at the plaque sites(78) manifested by impairment in motility and phagocytosis in mice with AD phenotype(79). This could lead to inactivation of the protective function of microglia to clear Aβ(80). Our observation that Aβ inhibits native PIEZO1 in human microglia supports the theory that the function of microglia is compromised in AD by completing the vicious circle exaggerating AD pathology. This, together with our observations that brain Aβ burden contributes to the expression of *PIEZO1* in microglia, indicates that dysfunction mechanisms triggering microglial malfunctions even before the diagnosed symptoms. Similar to the *in vivo* situation, *in vitro* Aβ preparations contain heterogenous Aβ species or blockage of PIEZO1 could be involved in the early ranging from monomeric to fibril forms. Even though our study does not specify which Aβ species mediate the inhibitory effects or through which exact mechanism, it is likely that Aβ can modulate cell membrane mechanical sensitivity to control PIEZO1-triggered Ca^2+^ influx. This could happen similarly as in human HEK293 cells rather than Aβ directly interacting with the PIEZO1 receptor(21). Thus, alleviating the Aβ-causing inhibitory effect on PIEZO1 offers a promising, early target for treatment paradigms. This view is further supported by the fact that the inhibition of PIEZO1 by the antagonists was toxic for iMGLs, although the toxicity could have also been a result of the unspecific off-target effects of the available inhibitors. Importantly, PIEZO1 activation in microglia provides a driver for multiple calcium-dependent mechanisms important for ‘waking up’ the protective function of these cells, including the immunological surveillance of the brain. This could be critically important for conditions such as AD where the proper function of microglia is compromised(81). This novel concept is in line with data showing that mechanosensitive stimuli, such as ultrasound, activates PIEZO1 channels (82) and leads to clearance of brain Aβ by microglia(83).

By using distinct mechanical cues and a selective molecular agonist we demonstrate the presence of abundant functional PIEZO1 mediated mechanotransduction in human iMGLs and mouse microglia. All the stimulation paradigms elicited prolonged functional responses lasting minutes and distinct from earlier observations of PIEZO1 mediated membrane currents in other cell types which were limited to ms range(84) due to the ion channel inactivation mechanism. The prolonged activity could arise from microglia-specific PIEZO1 organization, and in particular, the unusual organization of cytoskeletal structures in these mobile cells, and/or changes to membrane lipid organization shaping the global properties such as tension and fluidity. The PIEZO1 channel can switch permanently from a transient to a sustained gating mode by strong mechanical stimulation(85– 87) and by repeated stimulation(84,88). In particular, membrane cholesterol which is required for establishing a mature microglial phenotype (89), could affect the activity of PIEZO1 channel clusters(55,89,90).

Interestingly, we discovered that human iMGLs were highly sensitive to the PIEZO1 agonist Yoda1. Specific high sensitivity could stem from the species-specific structures of the PIEZO1 channel. Indeed, human and mouse share only 84 % similarity in the base pair sequence of the respective gene and 81% similarity in amino acids of the protein (https://blast.ncbi.nlm.nih.gov/). The semi-bell-shaped DRC of human cells contrasting to mouse microglia presented classical sigmoidal DRC could be due to specific PIEZO1 inactivation characteristics, analogous to similar contribution of desensitization to responses of capsaicin-activated Transient Receptor Potential Vanilloid (TRPV) receptors(91). Also differences in plasma membrane lipid and cholesterol composition could partly explain the differences between species since PIEZO1 activity appears to be concentrated in cholesterol-rich lipid raft domains(92,93) and human microglia, but not mouse microglia, show dysregulation of cholesterol homeostasis in sporadic AD models(94). In mice, global knock-out of *PIEZ*O is embryonically lethal, while in humans a PIEZO1 loss-of-function mutation has been reported to cause mainly a loss of lymphatic function(95), whereas a gain-of-function mutation results in a red blood cell dehydration(96) and is associated with protection from severe malaria in humans(97). All in all, the bell-shaped concentration dependence in iMGLs may have beneficial functional significance limiting potentially damaging cell Ca^2+^ overload at high Yoda1 concentrations(98).

Our calcium imaging experiments demonstrated that not all iMGLs responded similarly to Yoda1, suggesting that microglia cultures may have, similar to astrocytes(19), sub-populations that mediate the varieties in PIEZO1 responses. This indicates that bulk analysis of microglia could mask the altered pathways that are affected within specific subpopulations. Our reanalysis of single-cell transcriptomic datasets of AD patients(40) and DAM microglia in 5xFAD mice(41,42) showed apparently contradicting *PIEZO1* upregulation in a human AD-associated microglial subpopulation and, on the other hand, downregulation of *Piezo1* in mouse DAM. The original publications already stated that the human and mouse AD associated genes differ, and some genes can be even up- or downregulated to opposing directions, most likely reflecting species, brain region and/or disease stage differences. Despite the differences, the datasets demonstrated that *PIEZO1* positive microglia highly express some AD signature genes and correlate with some of the homeostatic genes that are downregulated in mouse DAM. This was in line with our pro- and anti-inflammatory metabolic profile induced by Yoda1 in human iMGLs suggesting that PIEZO1 microglia have a unique gene expression pattern reflecting their Aβ clearing function. To fully elucidate the role of microglia in PIEZO1-mediated clearance of Aβ at subpopulation level over the time-course of AD pathology in relation to the contribution of other cell types, a longitudinal single-cell analysis of subcellular clusters would be required.

Even though we are the first to focus on PIEZO1 in microglia specifically, we cannot rule out that Yoda1 may exert its beneficial actions also through other brain cell types such as astrocytes(19), oligodendrocytes(51), and neuronal stem cells(99) in AD(19,21,100). Being in line with a previous study of transgenic AD rats (19), our analysis shows a higher number of *PIEZO1* positive astrocytes in human AD brain. However, even though prior studies demonstrate that astrocytes express functional PIEZO1 channels(19,101), in our study Yoda1 did not alter GFAP immunoreactivity *in vivo*. In contrast, it specifically increased microglial IBA1 immunoreactivity and its localization around Aβ deposits. Moreover, not all astrocytes express PIEZO1 and only a portion of them upregulate it in response to certain pro-inflammatory stimuli and aging (19,51). Indeed, the lack of astrocytic contribution in our model could stem from young 5-month-old 5xFAD mice compared to 18-month-old rats in Velasco et al. 2018 study. These observations suggest that PIEZO1 channels may be implicated in both microglia and astrocytes in AD. However, microglial responses seem to precede and overhelm astrocytic contribution (19), following the paradigm that microglia are needed for astrocytic reactivation(102). Thus, we suggest that PIEZO1 in microglia plays a leading and mainly protective role in the context of AD pathology.

## CONCLUSIONS

In conclusion, we show the abundance and functional expression of PIEZO1 channels in microglia. Pharmacological stimulation of PIEZO1 may represent a unique approach to modulate a range of functions in human microglia. The observed reduction of brain Aβ in Yoda1-treated mice suggests that activation of PIEZO1 may be a potential treatment strategy for AD.

## METHODS

### Study design

Controlled laboratory experiments were used for studying AD-related cellular functions *in vitro* using human iPSC-derived cells, mouse primary and secondary cell cultures following to ISSCR Guidelines for Stem Cell Research. More complex *in vivo* tissue pathology was studied in an AD mouse model and WT control mice according to ARRIVE guidelines. For comparing to broader context both human and mouse transcriptional databases were utilized for in-depth analysis. The details of the measurement methods, subjects and units are described in the following sections. Sample size for animal experiments was determined by power analysis using a two-way ANOVA sample test for comparing means of two groups(22) using 0.80 power, 5% type I error rate, assumption of 1.5-fold change and 0.3 standard deviation to yield sample size of 6 animals per group. For functional *in vitro* analysis at least three technical replicates were used in each experiment. If the data indicated meaningful results, all data was repeated in three independent experiments and with three independent biological replicates. Results were presented as normalized to positive/negative control or vehicle to combine mean group values from multiple experiments in the same figure. For molecular analysis at least three independent samples were analyzed. Experiments were excluded and performed again in case there was no appropriate response to negative or positive controls. The weight of animals was measured before and after the treatments in case of need to exclude animals due to the weight loss. To minimize subjective bias, animals were randomized when allocated for groups using GraphPadQuickCalcs. Data analysis were blinded for the groups.

### Human iPSC-lines

Human iPSC-lines used in this study are listed in **Error! Reference source not found**.. All iPSCs lines were previously generated from skin biopsies collected from three female and seven male subjects at ages between 65 and 78-years and were characterized for their high quality, pluripotency, normal karyotypes and presence of APOE ε3/ε3 alleles(23– 27). In addition to healthy cell lines, ISOAD4 and ISOAD5 lines carrying AD-causing *PSEN1DE9* mutation were used(23). *PSEN1DE9* carriers were 48-years-old male with AD diagnosis and a presymptomatic 47-years old female and iPSC lines were edited by CRISPR/Cas9 system to create isogenic control lines where mutation comprising of 4.6 kb deletion in the exon 9 of the *PSEN1* gene(28) was corrected(23). In addition, a commercial BIONi010-C-2 line generated from a healthy 15-19 years-old male(29) and edited to have two APOE3/3 alleles was purchased from the European bank for Induced Pluripotent Stem cells (EBiSC) and produced and characterised by Bioneer (Copenhagen, Denmark). The producer reported that a DNA SNP array revealed no larger chromosomal aberrations, although an unsignificant duplication of 1,4Mbp was found on Chr22 in q11.23.

### Human induced pluripotent stem cell derived microglia-like (iMGL) cells

Prior to differentiation, iPSCs were maintained in serum-and feeder-free conditions in Essential 8 Medium (A15169-01 Gibco) with 0.5% penicillin-streptomycin (P/S, #15140122 Gibco) on growth factor reduced Matrigel (Corning) in a humidified incubator at 5% CO2 and 37°C. Cultures were passaged as small colonies with 0.5 mM EDTA (15575, Gibco) twice a week supplementing medium with 5 µM Rho-Associated Coil Kinase (ROCK) inhibitor Y-27632 (S1049, Selleckchem) for the first 24 hours. Passages under 50 were used and cultures were regularly tested for mycoplasma using a MycoAlert Kit (Lonza). Human iMGLs were differentiated as previously described(26). At differentiation day *D16*, cells were plated at different densities depending on the experiment and half of the medium was changed daily until *D21* to *D25*, when cells were treated for experiments.

### Human SV40 cell culture

Human immortalized microglial cell line SV-40 (T0251, Applied Biological Materials Inc.) originates from the Tardieu lab(30) and was obtained as a kind gift from Dora Brites, University of Lisbon, Portugal. The cells were maintained in DMEM + GlutaMAX 4.5 g/L glucose (10566016, Gibco) supplemented with 10% fetal bovine serum (FBS, Gibco) and 1% P/S in tissue culture T-25 flasks coated with 0.1 mg/mL collagen I (A10483-01, Gibco). For experiments, cells were seeded at 40,000 cells per cm^2^ density to produce ∼80% confluent cultures after 24 h for experiments.

### Primary murine microglial and astrocyte cell cultures

Primary murine microglial and astrocyte cultures were prepared from WT C57BL/6J neonatal mice on postnatal days P0–3 as described previously(31). Mice were sacrificed by decapitation and dissected brains were first dissociated mechanically and then enzymatically with 0.05% Trypsin-EDTA in DMEM/F-12 supplemented with 1% P/S and (all Gibco) into single cells. Cells per brain were seeded on a 15 cm tissue culture dish and were grown at 37°C, 5% CO2 for three weeks in DMEM/F-12, 10% FBS (Gibco) and 1% P/S to produce mixed glial culture. The astrocyte layer was detached with 30-minute incubtation in 0.8% Trypsin-EDTA and then microglia with 5 minutes in 0.25% Trypsin-EDTA. Both astrocytes and microglia were seeded in DMEM/F-12, 10% FBS and 1% P/S on Poly-D-lysine (PDL; P0899, Sigma) coated vessels one day prior to experiments.

### BV2 cell culture

The mouse microglial BV2 cell line was maintained in complete RPMI-1640 medium (Sigma), 1% GlutaMAX, 10% FBS and 5 μg/ml gentamicin (all Gibco) on 10 cm dishes for 2-10 passages. The cell line was confirmed to be negative for mycoplasma using MycoAlert Kit (Lonza).

### Trigeminal neuronal culture

Primary murine trigeminal neuron cultures were prepared from Wistar rats on P10 as previously described(32,33). Briefly, trigeminal ganglia were rapidly excised after decapitation and enzymatically dissociated in 0.25 mg/ml trypsin, 1 mg/ml collagenase I and 0.2 mg/ml DNAse (all Sigma) in F12 medium (Gibco) under continuous 850 rpm mixing at 37 °C for 15 min. The cells were seeded on 0.2 mg/ mL poly-l-lysine (PLL; Sigma) coated 24-mm glass coverslips (Thermo Fisher Menzel) and maintained at 37 °C, 5 % CO2 in F12 + GlutaMAX and 10% FBS (both Gibco) for 48 h prior to measurements.

### RT-qPCR gene expression analysis

Total RNA was extracted from brain tissues using the single-step acid guanidinium thiocyanate-phenol-chloroform method(34), and from iMGLs using mirVana™ miRNA Isolation Kit (Invitrogen) following manufacturer’s instructions. The concentration and purity of RNA were confirmed using a Nanodrop 2000 spectrophotometer (Thermo Fisher Scientific, Hudson, NH) with the 260 nm/280 nm ratio between 1.8 and 2. Total of 500 ng of RNA was reverse transcribed to cDNA using a Maxima reverse transcriptase with dNTP Mix (both Thermo Scientific). Gene expression levels were measured by RT-qPCR using StepOne Plus Real-Time PCR machine (Life Technologies) with FAM-labeled TaqMan probes (Thermo Scientific) with 40 cycles amplification. Probes are listed in (**Error! Reference source not found**.). Reactions were performed in triplicate with water and RNA controls to rule out contamination or genomic DNA. Relative gene expression was calculated as 2^(ΔΔCt) fold change compared to control group.

### Animals

Altogether, 5 WT C57BL/6J and 24 male transgenic 5XFAD mice (Jackson Laboratories, Bar Harbor, Maine, US) harboring the AD-linked Swedish (K670N/M671L), Florida (I716V), and London (V717I) mutations in human APP, and the M146L and L286V mutations in human PSEN1 genes were used. Mice were housed in a controlled environment within temperature 21 +/- 1 °C, humidity 50 +/-10%, light period from 7:00 a.m. to 7:00 p.m. and had ad libitum access to food and water. All mice were housed in groups of 3-7 animals in one cage with bedding and a piece of enrichment. Welfare of animals was daily monitored by animal facility personnel. No previous procedures were performed on any of the animals.

### *In vivo* treatments

To evaluate the effect of PIEZO activation*in vivo*, 5-month-old transgenic 5xFAD male mice (Jackson Laboratories, Bar Harbor, Maine, US) with fully developed Aβ pathology were used. For infusion, the animals were surgically implanted with intraventricular chronic cannulas as previously described(35). Briefly, anaesthesia was induced using 5% isoflurane (Attane vet®) in 30% O2/70% N2O and maintained at 1.8% isoflurane. The temperature of the animals was maintained at 37±0.5 °C using a thermostatically controlled heating blanket with a rectal probe (PanLab, Harvard Apparatus). A small hole approximately mm in diameter was drilled into the left hemisphere of the skull using coordinates: medial/lateral (m/l) +1.1 mm, anterior/posterior (a/p) −0.3 mm, dorsal/ventral (d/v) −2.0 mm. A chronic cannula (Cannula infusion system; Plastic1, Preclinical research components) was mounted into the left ventricle and fixed on the skull with dental mount. Skin was sutured, and mice were injected post-operatively with Temgesic for analgesia. Animals were placed in individual cages to recover for 24 h, followed by 5 µl cannula infusions of Yoda1 (0.25 µg/mouse/injection, Tocris) or equal volume of 1% DMSO in saline for a vehicle group once a day. The mice received total of 10 infusions over a period of 12 days, with 2-day break after the initial 5 infusions.

### Animal perfusion and tissue processing

The brain tissue samples were collected 6 hours after the last injection. The mice were terminally anesthetized using tribromoethanol (Avertin, Sigma-Aldrich) followed by transcardial perfusion with heparinized (2500 IU/L, LEO) 0.9% NaCl solution for four minutes at 19 ml/min speed. The brains were dissected out, cut midsagittal and the infused hemisphere was processed either for immunohistochemistry or for gene or protein analysis.

### Fixation and immunostainings

Immunohistochemistry was carried out as previously described(31). Left hemisphere was collected after perfusion and fixed in 4% paraformaldehyde (PFA, P6148, Sigma) in 0.1 M phosphate buffer (PB), pH 7.4 overnight at 4°C, followed by cryoprotection in 30% sucrose in PB for 48 hours at 4°C. The fixed hemispheres were snap-frozen on liquid nitrogen and cryosectioned using a cryotome (Leica CM1950) into 20-μm sagittal sections. Six consecutive sagittal brain sections at 400 μm intervals were stained from each mouse. *In vitro* cell cultures were fixed in 4% PFA in PBS, pH 7.4 for 20 min at RT or at +37°C for iMGLs. The cell membrane was permeabilized for 30 min in 0.4% Triton X-100 (Sigma) in PBS. Nonspecific binding was blocked in 10% NGS for 1 hour at RT. Sections and cell cultures were incubated with primary antibodies diluted in the application specific blocking solutions overnight at 4°C. All used antibodies are listed in **Error! Reference source not found**.3. Staining controls without primary antibodies were used to confirm staining specificity.

### Confocal and fluorescent microscopy

The confocal microscope images were obtained using 40x magnifications in a Zeiss Axio Observer inverted microscope equipped with LSM800 confocal module (Carl Zeiss Microimaging GmbH, Jena, Germany), fluorescent microscope images were received by Zeiss Axio Imager M.2 microscope equipped with Axiocam 506 mono CCD camera (Carl Zeiss, Oberkochen) using 10x magnification. DAPI and secondary antibodies were imaged with 405 nm (λex 353 nm/λem 465 nm), 488 nm (λex 495 nm/λem 519 nm), 568 nm (λex 543 nm/λem 567 nm) lasers, respectively. Images were captured using ZEN 2.3 Blue software (Carl Zeiss Microimaging GmbH) with constant conditions across negative controls and an experiment. The 3D super resolution z-stack images were taken at 0.5 µm Z-interval. Images were processed and exported using ZEN 2.3 Blue or ZEN 2.3 lite software’s (Carl Zeiss Microimaging GmbH). For figures confocal images were presented as maximum intensity projections or 3D images.

### Quantification of immunostainings

Quantitative analysis of each dataset was performed by two observers, who were blinded to the origin of the samples. For Aβ, GFAP, PIEZO1 and IBA1 staining optical fields of whole brain section were constructed by automatic stitching of individual image tiles using ZEN 2.3 Blue software. Quantification of hippocampal and cortical immunoreactivities from five consecutive slices was done using an unbiased, semi-automatic method using MatLab code (MathWorks, MatLab 2017b) and accuracy of analysis was confirmed by re-analyzing with FIJI (Wayne Rasband, National Institute of Health, USA, Collins, 2007; Schneider et al., 2012). The accumulation of microglia around Aβ deposits was quantified from confocal z-stack maximum intensity projection images. Projection images were divided to separate channel images and gaussian blur filter (sigma is 1) and threshold command were applied. The resulting binary images were further analyzed using particle command in FIJI software. The ratio of IBA1 staining per individual WO2 positive deposits was quantified in the ROIs manually circled around plaques from the center of the plaque in a way that it fit in the circleproperly through all the z-stack imagesand were divided based on the size of the deposits (small, medium and large)(36). Area size was detected automatically using Fiji tools.

### Calcium imaging

Calcium imaging was used for evaluating PIEZO1 evoked calcium influx in primary murine microglia, astrocytes and human iMGLs on PDL-coated round 5 mm glass coverslips (Thermo Scientific Menzel) at density of 7000-10000 cells/coverslip. The cells were loaded with 5 µM calcium-sensitive fluorescent dye Fluo-4AM (Direct Calcium Assay Kit, Invitrogen, USA) for 30 min at 37° C and washed for 15 min or 30 min at 37° C and then for 5 min at RT in basic solution (BS) containing at mM concentrations 5 KCl (Scharlau Chemicals), 152 NaCl, 1 MgCl2, 10 HEPES (all from VWR chemicals), 10 glucose (MP Biomedicals) and 2 CaCl2 (Merck KGaA) at pH 7.4. Coverslips were transferred in TILL photonics imaging system (TILLPhotonics GmbH, Germany) with constant BS perfusion. Fluorescence was visualized using monochromatic light source with ex/em 494/506 and 10x objective in Olympus IX-70 microscope (Tokyo, Japan) equipped with CCD camera (SensiCam, PCO imaging, Kelheim, Germany) using exposure time 100 ms and a frame per second sampling frequency.

To activate mechanosensitive channels, we used: 1) fluid pressure pulse; 2) hypo-osmotic solution (HOS); 3) a selective PIEZO1 agonist Yoda1 (5586, Tocris). For pressure pulse stimulation, a Picospritzer III (Parker Hannifin, USA) connected with pressure supplier source (N2 tank) was used. Stimulation was produced by 30 psi (500 ms) puff of BS from the glass pipette filled with BS (resistance 1.5 Mom) positioned next to the target cell at equal distance for each measurement. After 1-minute recovery, 0.1 µM Yoda1 was applied for 1 min to the same cell. For osmotic and chemical stimulation, HOS and Yoda1 were applied via fast perfusion system (Rapid Solution Changer RSC-200, BioLogic, Grenoble, France) at 2.5 ml/sec flow speed (∼30 ms solution exchange time) for 1-2 min. For HOS, NaCl was reduced in BS to 81 mM to get 40% lower osmotic value of 200 mOsm/kg instead of 320 mOsm/kg of isotonic BS. For vehicle control, DMSO in correspondent concentration was added to perfusion BS across experiments. To study PIEZO1 inhibition, cells were incubated for 1 h with 5 µM GsMTx4 (Tocris) or 15 and 30 min with 1 µM soluble Aβ1-42 (Bachem) prior to Yoda1 application. To quantify the amplitude of calcium transients, the ratio between the peak amplitude of calcium responses were normalized after subtracting a baseline. Data were pre-analyzed offline using ImageJ (Rasband, W.S., ImageJ, U. S. National Institutes of Health, Bethesda, Maryland, USA, https://imagej.nih.gov/ij/) and results were obtained via Origin 2019b (OriginLab, Northampton, Massachusetts, USA).

### Real time Cytotox Green cytotoxicity assay

Cytotox Green assay (4633, Essen Bioscience) was used for *in vitro* iMGLs to obtain cell death overtime in Incucyte S3 Live-Cell Analysis System (4647, Essen Bioscience, Ann Arbor, MI, USA). Matured iMGLs at density of 15 000 cells/well on 96-well plates were treated with 0.3-50 µM Yoda1 (Tocris) or equal volume of DMSO, both with 250 nM Cytotox Green reagent. As a positive control, 200 µM 1-Methyl-4-phenylpyridinium iodide (D048, MPP+, Sigma) was applied to induce cell death over 60 hours. Cells were live-imaged in IncuCyte S3 at 37 °C and 5% CO2 every 3 hours for 3 days. One to two images/well were captured using 10x objective for phase-contrast to obtain confluency and for green fluorescence to obtain intensity of reagent binding to DNA of dying cells with compromised membrane. IncuCyte S3 Software (2019B) with constant settings across conditions was used for quantification. Green integrated intensity as was thresholded from unspecific background fluorescence with a top-hat thresholding to get green green calibrated unit (GCU). GCU per image was divided by confluence area per image at each time point. Data from independent iMGL batches and experiments was combined by normalizing to control group.

### LDH assay

CyQUANT™ LDH Cytotoxicity Assay Kit (Invitrogen, C20301) according to manufacturer’s instructions was used for measuring the release of cytosolic lactate dehydrogenase (LDH) enzyme into the cell culture media as indicative of compromised cell membrane aka cytotoxicity. The cells were seeded and treated similarly as for cytotox green assay and the medium was collected after 24 h. In the assay, the extracellular LDH catalyzes the conversion of lactate to pyruvate via NAD+ reduction to NADH accompanied with a red formazan formation proportional to the amount of LDH. The absorbances at 490 nm and 680 nm were measured using Victor Wallace plate reader and LDH activity determined by subtracting the background absorbance and spontaneous LDH activity and then normalizing to the positive control of 2 % Triton X-100.

### Mitostress

Mitostress assay was performed using a Seahorse XFe96 Extracellular Flux Analyzer (Agilent) to measure OCR and ECAR in realtime following the manufacturer instructions. IMGLs were seeded as a monolayer with 50 000 cells/well in a XF96-well Seahorse microplate (Agilent). Cells were washed and equilibrated in the XF Assay modified DMEM medium for 1 h at 37°C non-CO_2_ incubator. The levels of OCR and ECAR were determined in response to the sequential addition of oligomycin, FCCP and rotenone/antimycin A (all 1 mM and from Sigma). Non-mitochondrial respiration was determined as the minimum rate measurement after addition of Rotenone/Antimycin A. Basal mitochondrial respiration was calculated by subtracting non-mitochondrial respiration from the last measurement before addition of oligomycin. Spare capacity was calculated by subtracting basal respiration from the maximum rate measurement after addition of FCCP. ATP production was determined by subtracting the minimum rate after adding oligomycin from the last previous measurement before addition of oligomycin. The parameters were obtained using the Agilent Report Generator in Wave software.

### Scratch wound migration assay

Chemokinesis of iMGLs was analyzed using IncuCyte® scratch wound cell migration assay (Sartorius) with real-time visualization according to manufacturer’s instructions. To achieve confluent monolayers, iMGLs were seeded at density of 30 000 cells/well on PDL coated ImageLock 96-well plates. The cells were pretreated with Yoda1, equal volume of DMSO as vehicle. After 22 h medium was changed to make the adherent cells more prone to detach. After 2 hours, the monolayers were wounded using a 96-well WoundMaker (Sartorius) to create coincident 700-800 μm wide wounds to each well. The medium was again changed to remove detached cells. Location-matched 10× phase-contrast images were captured once per hour for 24 hours using the scratch wound mode and IncuCyte S3 scratch wound cell migration software module (version 2019B) was used for quantifying relative wound density using default settings. The percentage of relative wound density compared to vehicle group was calculated by normalizing to maximum value of vehicle.

### Phagocytosis assays

Phagocytosis of iMGLs was evaluated with fluorescent HiLyte Fluor 488 Aβ1-42 (AnaSpec) and by pHrodo BioParticle (Invitrogen) phagocytosis assays using IncuCyte S3 live-cell imaging as previously described(26). Cells were seeded similarly as for Cytotox assay and were pretreated for 24 h in FBS-omitted PM medium. After 24 h, 1 µM HiLyte Aβ1-42 or 62.5µg/ml pHrodo bioparticles were applied in Opti-MEM (Gibco) on the top of old medium and one to two 20× images/well were taken every 30 minutes for 6 hours using the phase-contrast and green fluorescence to obtain time curves in IncucCyte. Background fluorescence of non-phagocytosed beads/ HiLyte Aβ1-42 was removed using top-hat thresholding similarly as for cytotox assay in IncuCyte S3 software (version 2019B). For pHrodo bioparticles that were visible in brightfield images, GCU was normalized to confluence captured before addition of pHrodo particles, while HiLyte Aβ1-42 was normalizaed to confluence at each timepoint.

### Lysosomal activity

Lysosomal activity was measured for the BV2 cell line and iMGLs using the LysoViewTM 540 reagent (Biotium, 70061) and Incucyte S3 Live-Cell imaging. BV2 cells were seeded for the assay at density of 10 000 cells/well on PDL (Sigma, P0899) coated 96-well plates and iMGLs as for cytotox assay. LysoView reagent was added, and the first image taken (T=0) was used to evaluate the basal lysosomal activity per well. Cells were treated with Yoda or equal volume of DMSO as the vehicle control. Two 20x images per well were captured every 3 hours for 24 hours for phase-contrast confluence and for red fluorescence to measure accumulating intensity of the LysoViewTM 540 reagent in lysosomes. Red fluorescence was separated from background with a top-hat thresholding and red integrated intensity per image was divided by confluence area per image at each time point. The data were normalized to the T=0 to avoid the bias due to different fluorescence intensity between wells and then compared to maximum values of positive control or vehicle.

### Analysis of RNA-seq datasets

To quantitatively compare PIEZO1 gene expression in different human cell types, RNA-seq data from two published datasets of human iPSC-microglia with GEO accession numbers GSE89189(37) and GSE135707(26) were implemented as previously described(26). Both datasets were created from total RNA with a RIN score >9 using TruSeq mRNA stranded protocols to obtain poly-A mRNA and were sequenced using Illumina HiSeq instuments. In both cases RNA-seq reads were mapped to the hg38 reference genome. To be able to compare datasets with different origins and sequenced as single-end 50 bp reads or paired-end 100 bp reads the raw data were preprocessed. Briefly, raw data were log2-transformed and quantile normalized using the function “normalize.quantiles” of the package preprocessCore(38). Subsequently, data were batch-corrected to remove the bias due to the technical variability of the two studies. Batch correction was performed using the function “removeBatchEffect” of the limma package.

### Clustering analysis of snRNA- and scRNA-seq datasets

Seurat(39) was used for the analysis of the Grubman et al. snRNA-seq(40), Keren-Shaul et al. scRNA-seq(41) and Zhou et al. snRNA-seq (42) datasets. For Zhou et al. data we removed the cells identified as multiplets by performing the scds method(43). Following Seurat and to improve the precision of the scRNA/snRNA analysis, a gene was included if present in at least 3 cells, while a cell was included if it expressed at least 200 genes(44). The cells which had a percentage of mitochondrial genes greater than 5% were considered as dying cells and were filtered out(45). A further filtering was done following the original authors of the data based on the number of unique molecular identifiers (UMI) and of genes per cell. After this processing, data were normalized to reduce technical differences(46), scaled(44) and clustered. PCA and JackStraw procedures were applied based on the 2000 most variables and the number of PCA components was then chosen at the ‘elbow’ of the plot. The first 20 PCA components ensured the best separation of the cells and clusters have been detected with a resolution equal to 0.32 for the Grubman et al. dataset, while 0.2 for Keren-Shaul and 0.06 for Zhou data. Finally, the cluster annotation in specific cell-types was addressed using scCATCH(47) for the human dataset, while using gene set variation analysis (GSVA)(48), for the mouse ones. The latter has been applied as defined by J. Javier et al.(49) and the tested gene sets have been composed by Keren-Shaul and Zhou cell-type specific most expressed genes published in the supplementary material of the original papers.

After the cell clustering, the gene statistics in the clusters were analyzeed. We determined the ratio of frequency and of expression to understand the presence and the level of expression of each gene with respect the others (e.g. frequency ratio of 0.6 means that the gene is expressed in more cells than 60% of the other genes in the same cell-type). To conclude, we summarized the information retrieved for each gene. We defined a continuous numeric score, from the minimum value of 0 to the maximum of 2, based on the sum of the ratio of frequency and the ratio of expression. For example, the gene which in a cell-type is both the most frequent and the most expressed with respect the other genes get a score equal to 2, while 0 otherwise. We then performed a correlation analysis between the DAM markers (Keren-Shaul’s et al. genes associated to AD-specific microglia) and *Piezo1*. We determined how much the DAM markers were correlating with their expression to *Piezo1* expression in the mouse microglia subpopulations. In case of the Keren-Shaul’s et al data, we considered the gene expressions in the Microglia 1, 2 3. In case of the Zhou’s et al. data, we considered the gene expressions in the WT_5XFAD (1) and in the microglia 0 and 2. We selected the DAM markers that got a high correlation with *Piezo1* and that were present in at least the 70% of the *Piezo1* cells. We kept only the DAM markers satisfying the criteria in both the two mouse datasets. All the analysis can be replicated with the R>=3.6 scripts available at the following link: https://github.com/LucaGiudice/Microglia-AD-PIEZO1.

### Statistical analysis

All quantitative assessment was performed in a blinded manner and based on power calculation wherever it was possible. All cell culture experiments were independently repeated at least three times. For iMGLs each experiment was performed in an independent batch of cells differentiated from iPSCs. Based on the type and distribution of data populations (examined with Shapiro-Wilks W test) appropriate statistical tests were applied: for two independent groups, two-tailed unpaired t-test or non-parametric Mann Whitney t-test, for two dependent groups of data Wilcoxon signed-rank test, for multiple comparisons one-way ANOVA (with Tukey’s or Bonferroni’s post hoc tests) or Kruskal-Wallis test was used. The analysis was performed with the GraphPad Prism v.8 for Windows (GraphPad Software, La Jolla California USA, www.graphpad.com) and Origin2019b (OriginLab, for Ca imaging experiments). Outliers were removed using extreme studentized deviate Grubbs’ test with 0.05 significance level. All data is represented as median or mean ± SEM, unless otherwise stated. Differences with p < 0.05 were considered significant. Statistically significant differences were set at *p < 0.05 and **p < 0.01 and ***p < 0.001.

## DECLARATIONS

### Ethics approval and consent to participate

Human iPSC-lines were used compliant with the Decleration of Helsinki of the WMA, 1964. Written informed consent was obtained from all the participants. MBE2968, MBE2960 and TOB0644 were generated upon approval from the human research ethics committees of the Royal Victorian Eye and Ear Hospital (11/1031H), University of Melbourne (1545394), University of Tasmania (H0014124), with the requirements of the National Health & Medical Research Council of Australia and conformed with the Declarations of Helsinki(24,26,103). Ctrl 8, LL190, MAD1, MAD6, MAD8, ISOAD4 and ISOAD5 lines were generated upon approval from the committee on Research Ethics of Northern Savo Hospital district (license no. 123/2016)(23). The other lines were obtained commercially.

All animal experiments were performed in accordance with the EU Directive 2010/63/EU on the protection of animals used for scientific purposes and were approved and monitored by the Animal Experiment Board in Finland. All efforts were made to minimize the number of animals used and the wellbeing of animals was considered in all steps.

## Consent for publication

Not applicable.

## Availability of data and materials

All data, code, and materials used in the analysis are available from the corresponding author upon reasonable request or accessed at https://github.com/LucaGiudice/Microglia-AD-PIEZO1.

## Authors’ contributions

The contributions are visualized in Fig S6 based on CRediT Taxonomy(104). Conceptualization: HJ, VS, YI, RG RashidG, TM. Methodology: HJ, VS, YI, AS, IFU, NM, PK, KK, SL, DH, AP. Investigation: HJ, VS, YI, AS, IFU, MGB, NK, NM, SO, IF, IS, IB, MG, PK. Formal analysis: HJ, VS, YI, AS, LG, IFU, NK, PK. Validation: HJ, VS, YI, AS, LG, IFU, NK, SO, SL, DH, AP, RashidG. Software: GL, HJ, VS, YI, NK, RashidG. Data curation: HJ, VS, YI, AS, LG, IFU, NK. Visualization: HJ, VS, YI, AS, LG, IFU, RG RashidG, TM. Funding acquisition: HJ, SL, JK, KMK, AP, PK, RG, RashidG, TM. Resources: JK, KMK, AP, RashidG, RG, TM. Project administration: HJ, VS, YI, PK, RG RashidG, TM. Supervision: HJ, YI, JK, PK, RG RashidG, TM. Writing – original draft: HJ, VS, YI, AS, LG, IFU, PK, RG, RashidG, TM. Writing – review & editing: HJ, VS, YI, AS, LG, IFU, KMK, AP, RashidG, RG, TM.

## Acknowledgements

We thank FIMM Digital Microscopy and Molecular Pathology Unit supported by HiLIFE and Biocenter Finland for Zeiss Airyscan super-resolution imaging of histological stainings of tissue samples. We thank Mrs Mirka Tikkanen from University of Eastern Finland for the assistance with immunohistochemistry, Iveta Stanova, Feroze Fazaludeen and Ilkka Fagerlund for their help in stem cell maintanance, Dzhessi Rait for help with cannulation of animals, and Nikolay Naumenko and Pasi Tavi for lending of equipment for the mechanical microglia stimulation. This work was carried out with the support of UEF Cell and Tissue Imaging Unit, andcomputational analyses were performed on servers provided by UEF Bioinformatics Center, University of Eastern Finland.

## LIST OF ABBREVIATIONS

AD: Alzheimer’s disease
ADP: Adenosine diphosphate
ANOVA: Analysis of variance
APOC1: Apolipoprotein C1
APOE: Apolipoprotein E
APP: Amyloid-beta precursor protein
ARRIVE: Animal Research: Reporting of In Vivo Experiments
ATP: Adenosine triphosphate
AUC: Area under curve
Aβ: Amyloid beta
BF: Bright field
Cas9: CRISPR-associated protein 9
CCD: Charge-coupled device
CD14: Cluster of differentiation 14
CD14M: CD14+ monocyte
cDNA: Complementary DNA
Cox6a1: Cytochrome c oxidase subunit 6A1
CRISPR: Clustered Regularly Interspaced Short Palindromic Repeats
Ctsb: Cathepsin B
Ctsd: Cathepsin D
Ctsl: Cathepsin L
Ctsz: Cathepsin z
CX3CR1: CX3C chemokine receptor 1
DAM: Disease associated microglia
DAPI: 4′,6-diamidino-2-phenylindole
DC: Dendritic cell
DEG: Differentially expressed gene
DMEM: Dulbecco’s Modified Eagle Medium
DMSO: Dimethyl sulfoxide
DNA: Deoxyribonucleic acid
DNAse: Deoxyribonuclease
dNTP: Deoxynucleoside triphosphate
DRC: Dose-response curves
EBiSC: European bank for Induced Pluripotent Stem cells
ECAR: Extracellular acidification
EDTA: Ethylenediaminetetraacetic acid
EOAD: Early-onset Alzheimer’s disease
Expr.: Expression level
F12: Nutrient Mixture F-12
FBS: Fetal bovine serum
FCCP: Carbonyl cyanide-p-trifluoromethoxyphenylhydrazon
FIMM: Institute for Molecular Medicine Finland
FPKM: Fragments Per Kilobase of transcript per Million mapped reads
Freq.: frequency
FRMD4A: FERM domain containing 4A
Fth1: Ferritin heavy chain 1
GCU: Green calibrated Unit
GFAP: Glial fibrillary acidic protein
Glul: Glutamate-ammonia ligase
GSVA: Gene set variation analysis
GWAS: Genome-wide association study
HEK293: Human embryonic kidney 293 cells
HiLiFE: Helsinki Institute of Life Science
HOS: Hypo-osmotic solution
IBA1: Ionized calcium binding adaptor molecule 1
IFNγ: Interferon Gamma
IHC: Immunocytochemistry
iHPC: Induced hematopoietic progenitor cells
IL-13: Interleukin 13
IL-4: Interleukin 4
iMGL: iPSC-derived microglia-like cells
iPSC: Induced pluripotent stem cell
ISSCR: The International Society for Stem Cell Research
LDH: Lactate dehydrogenase
LINGO1: Leucine rich repeat and Ig domain containing 1
LOAD: Late-onset Alzheimer’s disease
LPS: Lipopolysaccharide
Malat1: Metastasis associated lung adenocarcinoma transcript 1
Marcks: Myristoylated alanine rich protein kinase C substrate
MG: Microglia
miRNA: MicroRNA
MPP+: 1-Methyl-4-phenylpyridinium iodide
MT-ATP6: Mitochondrially encoded ATP synthase membrane subunit 6
MT-CO2: Mitochondrially encoded cytochrome c oxidase II MT-
CO3: Mitochondrially encoded cytochrome c oxidase III
MT-CYB: Mitochondrially encoded cytochrome b
MT-ND2: Mitochondrially encoded NADH:ubiquinone oxidoreductase core subunit 2
MT-ND4: Mitochondrially encoded NADH:ubiquinone oxidoreductase core subunit 4
NGS: Normal goat serum
Npm1: Nucleophosmin 1
OCR: Oxygen consumption rate
OXPHOS: Oxidative phosphorylation
P/S: Penicillin Streptomycin solution
P2RY12: Purinergic Receptor P2Y12
PB: Phosphate buffer
PBS: Phosphate-buffered saline
PC: Positive control
PCA: Principal component analysis
PCR: Polymerase chain reaction
PCT: Patent Cooperation Treaty
PDL: Poly-d-lysine
PFA: Paraformaldehyde
PIEZO1: Piezo Type Mechanosensitive Ion Channel Component 1
PLL: Poly-L-lysine
PSEN1: Presenilin 1
RIN: RNA integrity number
RNA: Ribonucleic acid
ROCK: Rho-associated protein kinase inhibitor
RPKM: Reads Per Kilobase of transcript per Million mapped reads
Rpl22l1: Ribosomal protein L22 like 1
Rpl35a: Ribosomal protein L35a
Rpl5: Ribosomal protein L5
Rplp2: Ribosomal protein lateral stalk subunit P2
Rps11: Ribosomal protein s11
Rpsa: Ribosomal protein SA
RT-qPCR: Rreal-time quantitative polymerase chain reaction
scRNA-seq: Single-cell RNA sequencing
SEM: Standard error of mean
Serinc3: Serine incorporator 3
SNP: Single nucleotide polymorphism
snRNA-seq: Single nucleus RNA sequencing
SNX6: Sorting nexin 6
TMEM119: Transmembrane Protein 119
Tmem173: Stimulator of interferon response cGAMP interactor 1
TREM2: Triggering Receptor Expressed On Myeloid Cells 2
UEF: University of Eastern Finland
UMAP: Uniform Manifold Approximation and Projection
UMI: Unique Molecular Identifier
VEH: Vehicle
WMA: World Medical Association
WT: Wild type

## SUPPLEMENTARY INFORMATION

Additional file 1: Supplementary figures and tables.

Fig. S1. *PIEZO1* and *PIEZO2* expression in human and mouse RNA-seq dataset and staining controls for hiMGL immunostaining.

Fig. S2. Activation of PIEZO1 orchestrates immune response of human iMGLs.

Fig. S3. No differences in PIEZO1 expression in bulk brain tissue between WT and 5xFAD mice.

Fig. S4. PIEZO1, microglia, astrocyte and Aβ plaque stainings in 5xFAD hippocampi.

Fig. S5. *PIEZO1* gene expression in published AD-related RNA datasets by our analysis.

Fig. S6. A diagram visualizing author contribution.

Fig. S7. A graphical abstract summarizing the main finding of the paper.

Table S1. A correlation data for *Piezo1* and the DAM signature genes in microglial subpopulations in Keren-Shaul et al. 2017 dataset(41) (GSE98969).

Table S2. A correlation data for *Piezo1* and the DAM signature genes in microglial subpopulations in Zhou et al. 2021 dataset(42) (GSE140511).

Table S3. Differentially expressed genes (DEGs) specific for m1-subcluster in snRNA Grubman et al. 2019 dataset (40)(GSE138852).

## REFERENCES AND NOTES

1. Hammond TR, Robinton D, Stevens B. Microglia and the Brain: Complementary Partners in Development and Disease. Vol. 34, Annual Review of Cell and Developmental Biology. 2018.

2. Malm TM, Jay TR, Landreth GE. The Evolving Biology of Microglia in Alzheimer’s Disease. Vol. 12, Neurotherapeutics. Springer New York LLC; 2015. p. 81–93.

3. Mawuenyega KG, Sigurdson W, Ovod V, Munsell L, Kasten T, Morris JC, et al. Decreased clearance of CNS β-amyloid in Alzheimer’s disease. Science (80-). 2010 Dec 24 ;330(6012):1774. /pmc/articles/PMC3073454/

4. Wildsmith KR, Holley M, Savage JC, Skerrett R, Landreth GE. Evidence for impaired amyloid β clearance in Alzheimer’s disease. Vol. 5, Alzheimer’s Research and Therapy. BioMed Central; 2013. p. 33. http://alzres.biomedcentral.com/articles/10.1186/alzrt187

5. Hickman SE, Allison EK, El Khoury J. Microglial dysfunction and defective β-amyloid clearance pathways in aging Alzheimer’s disease mice. J Neurosci. 2008;28(33):8354–60.

6. Linnartz-Gerlac CB; VLADB. Engineered stem cell-derived microglia as therapeutic vehicle for experimental autoimmune encephalomyelitis - NONE. Gene Therapy. 2013. p. 797–806.

7. Moeendarbary E, Weber IP, Sheridan GK, Koser DE, Soleman S, Haenzi B, et al. The soft mechanical signature of glial scars in the central nervous system. Nat Commun. 2017 Mar;8.

8. ElSheikh M, Arani A, Perry A, Boeve BF, Meyer FB, Savica R, et al. MR elastography demonstrates unique regional brain stiffness patterns in dementias. Am J Roentgenol. 2017 Aug;209(2):403–8.

9. Murphy MC, Jones DT, Jack CR, Glaser KJ, Senjem ML, Manduca A, et al. Regional brain stiffness changes across the Alzheimer’s disease spectrum. NeuroImage Clin. 2016;10:283–90.

10. Smith JF, Knowles TPJ, Dobson CM, Macphee CE, Welland ME. Characterization of the nanoscale properties of individual amyloid fibrils. 2006.

11. Bollmann L, Koser DE, Shahapure R, Gautier HOB, Holzapfel GA, Scarcelli G, et al. Microglia mechanics: immune activation alters traction forces and durotaxis. Front Cell Neurosci. 2015 Sep 23 ;9(September):363. http://journal.frontiersin.org/Article/10.3389/fncel.2015.00363/abstract

12. Moshayedi P, Ng G, Kwok JCF, Yeo GSH, Bryant CE, Fawcett JW, et al. The relationship between glial cell mechanosensitivity and foreign body reactions in the central nervous system. Biomaterials. 2014 Apr;35(13):3919–25.

13. Condello C, Yuan P, Schain A, Grutzendler J. Microglia constitute a barrier that prevents neurotoxic protofibrillar Aβ42 hotspots around plaques. Nat Commun. 2015 May 29;6(1):6176. http://www.nature.com/authors/editorial_policies/license.html#terms

14. Coste B, Xiao B, Santos JS, Syeda R, Grandl JJ, Spencer KS, et al. Piezo proteins are pore-forming subunits of mechanically activated channels. Nature. 2012 Mar 19;483(7388):176–81. http://www.nature.com/articles/nature10812

15. Coste B, Mathur J, Schmidt M, Earley TJ, Ranade S, Petrus MJ, et al. Piezo1 and Piezo2 are essential components of distinct mechanically activated cation channels. Science (80-). 2010 Oct 1;330(6000):55–60.

16. Bron R, Wood RJ, Brock JA, Ivanusic JJ. Piezo2 expression in corneal afferent neurons. J Comp Neurol. 2014;522(13):2967–79.

17. Eijkelkamp N, Linley JE, Torres JM, Bee L, Dickenson AH, Gringhuis M, et al. A role for Piezo2 in EPAC1-dependent mechanical allodynia. Nat Commun. 2013;4.

18. Kim SE, Coste B, Chadha A, Cook B, Jolla L, Diego S. The role of Drosophila Piezo in mechanical nociception. 2012;483(7388):209– 12.

19. Velasco-Estevez M, Mampay M, Boutin H, Chaney A, Warn P, Sharp A, et al. Infection Augments Expression of Mechanosensing Piezo1 Channels in Amyloid Plaque-Reactive Astrocytes. Front Aging Neurosci. 2018 Oct 22;10(OCT):332. https://www.frontiersin.org/article/10.3389/fnagi.2018.00332/full

20. Wu J, Lewis AH, Grandl J. Touch, Tension, and Transduction - The Function and Regulation of Piezo Ion Channels. Trends Biochem Sci. 2017 Jan;42(1):57–71.

21. Maneshi MM, Ziegler L, Sachs F, Hua SZ, Gottlieb PA. Enantiomeric Aβ peptides inhibit the fluid shear stress response of PIEZO1. Sci Rep. 2018 Dec;8(1).

22. Chow S, Shao J, Wang H. Sample Size Calculations in Clinical Resarch. 2nd ed. Chapman & Hall/CRC Biostatistics Series; 2008. 58 p.

23. Oksanen M, Petersen AJ, Naumenko N, Puttonen K, Lehtonen Š, Gubert Olivé M, et al. PSEN1 mutant iPSC-derived model reveals severe astrocyte pathology in Alzheimer’s disease. Stem Cell Reports. 2017 Dec 12;9(6):1885–97.

24. Muñoz SS, Engel M, Balez R, Do-Ha D, Cabral-da-Silva MC, Hernández D, et al. A Simple Differentiation Protocol for Generation of Induced Pluripotent Stem Cell-Derived Basal Forebrain-Like Cholinergic Neurons for Alzheimer’s Disease and Frontotemporal Dementia Disease Modeling. Cells. 2020 Sep;9(9).

25. Guneykaya D, Ivanov A, Hernandez DP, Haage V, Wojtas B, Meyer N, et al. Transcriptional and Translational Differences of Microglia from Male and Female Brains. Cell Rep. 2018 Sep 4;24(10):2773-2783.e6. https://pubmed.ncbi.nlm.nih.gov/30184509/

26. Konttinen H, Cabral-da-Silva M e. C, Ohtonen S, Wojciechowski S, Shakirzyanova A, Caligola S, et al. PSEN1ΔE9, APPswe, and APOE4 Confer Disparate Phenotypes in Human iPSC-Derived Microglia. Stem Cell Reports. 2019 Oct 8;13(4):669–83.

27. Fagerlund I, Dougalis A, Shakirzyanova A, Gómez-Budia M, Pelkonen A, Konttinen H, et al. Microglia-like Cells Promote Neuronal Functions in Cerebral Organoids. Cells 2022, Vol 11, Page 124. 2021 Dec 30;11(1):124. https://www.mdpi.com/2073-4409/11/1/124/htm

28. Crook R, Verkkoniemi A, Perez-Tur J, Mehta N, Baker M, Houlden H, et al. A variant of Alzheimer’s disease with spastic paraparesis and unusual plaques due to deletion of exon 9 of presenilin 1. Nat Med. 1998;4(4):452–5.

29. Rasmussen MA, Holst B, Tümer Z, Johnsen MG, Zhou S, Stummann TC, et al. Transient p53 suppression increases reprogramming of human fibroblasts without affecting apoptosis and DNA damage. Stem Cell Reports. 2014 Sep;3(3):404–13.

30. Janabi N, Peudenier S, Héron B, Ng KH, Tardieu M. Establishment of human microglial cell lines after transfection of primary cultures of embryonic microglial cells with the SV40 large T antigen. Neurosci Lett. 1995 Aug;195(2):105–8.

31. Malm T, Mariani M, Donovan LJ, Neilson L, Landreth GE. Activation of the nuclear receptor PPARδ is neuroprotective in a transgenic mouse model of Alzheimer’s disease through inhibition of inflammation. J Neuroinflammation. 2015 Jan;12:7. http://www.pubmedcentral.nih.gov/articlerender.fcgi?artid=4310027&tool=pmcentrez&rendertype=abstract

32. Yegutkin GG, Guerrero-Toro C, Kilinc E, Koroleva K, Ishchenko Y, Abushik P, et al. Nucleotide homeostasis and purinergic nociceptive signaling in rat meninges in migraine-like conditions. Purinergic Signal. 2016;12(3):561–74.

33. Ceruti S, Fumagalli M, Villa G, Verderio C, Abbracchio MP. Purinoceptor-mediated calcium signaling in primary neuron-glia trigeminal cultures. Cell Calcium. 2008;43(6):576–90.

34. Chomczynski P, Sacchi N. Single-step method of RNA isolation by acid guanidinium thiocyanate-phenol-chloroform extraction. Anal Biochem. 1987 Apr;162(1):156–9.

35. DeVos SL, Miller TM. Direct intraventricular delivery of drugs to the rodent central nervous system. J Vis Exp. 2013;(75):50326.

36. Bolmont T, Haiss F, Eicke D, Radde R, Mathis CA, Klunk WE, et al. Dynamics of the microglial/amyloid interaction indicate a role in plaque maintenance. J Neurosci. 2008 Apr 16;28(16):4283–92.

37. Abud EM, Ramirez RN, Martinez ES, Healy LM, Nguyen CHH, Newman SA, et al. iPSC-Derived Human Microglia-like Cells to Study Neurological Diseases. Neuron. 2017 Apr 19;94(2):278-293.e9. http://www.sciencedirect.com/science/article/pii/S0896627317302866

38. Bolstad B, Irizarry R, Astrand M, Speed T. A Comparison of Normalization Methods for High Density Oligonucleotide Array Data Based on Variance and Bias - PubMed. Bioinformatics. 2003;19(2):185–93. https://pubmed.ncbi.nlm.nih.gov/12538238

39. Butler A, Hoffman P, Smibert P, Papalexi E, Satija R. Integrating single-cell transcriptomic data across different conditions, technologies, and species. Nat Biotechnol. 2018 Jun 1;36(5):411–20. 40

40. Grubman A, Chew G, Ouyang JF, Sun G, Choo XY, McLean C, et al. A single-cell atlas of entorhinal cortex from individuals with Alzheimer’s disease reveals cell-type-specific gene expression regulation. Nat Neurosci. 2019 Dec 1;22(12):2087–97.

41. Keren-Shaul H, Spinrad A, Weiner A, Matcovitch-Natan O, Dvir-Szternfeld R, Ulland TK, et al. A Unique Microglia Type Associated with Restricting Development of Alzheimer’s Disease. Cell. 2017 Jun 15;169(7):1276-1290.e17.

42. Zhou Y, Song WM, Andhey PS, Swain A, Levy T, Miller KR, et al. Human and mouse single-nucleus transcriptomics reveal TREM2-dependent and TREM2-independent cellular responses in Alzheimer’s disease. Nat Med. 2020 Jan 13;26(1):131–42. http://www.nature.com/articles/s41591-019-0695-9

43. Bais AS, Kostka D. Scds: Computational annotation of doublets in single-cell RNA sequencing data. Bioinformatics. 2020 Feb 15;36(4):1150–8. https://github.com/JonathanShor/DoubletDetection,

44. Wu Y, Zhang K. Tools for the analysis of high-dimensional single-cell RNA sequencing data. Vol. 16, Nature Reviews Nephrology. Nature Research; 2020. p. 408–21.

45. Potter SS. Single-cell RNA sequencing for the study of development, physiology and disease. Vol. 14, Nature Reviews Nephrology. Nature Publishing Group; 2018. p. 479–92.

46. Vallejos CA, Risso D, Scialdone A, Dudoit S, Marioni JC. Normalizing single-cell RNA sequencing data: Challenges and opportunities. Vol. 14, Nature Methods. Nature Publishing Group; 2017. p. 565–71.

47. Shao X, Liao J, Lu X, Xue R, Ai N, Fan X. scCATCH: Automatic Annotation on Cell Types of Clusters from Single-Cell RNA Sequencing Data. iScience. 2020 Mar;23(3):100882.

48. Hänzelmann S, Castelo R, Guinney J. GSVA: Gene set variation analysis for microarray and RNA-Seq data. BMC Bioinformatics. 2013 Jan 16;14. https://pubmed.ncbi.nlm.nih.gov/23323831/

49. Diaz-Mejia JJ, Meng EC, Pico AR, MacParland SA, Ketela T, Pugh TJ, et al. Evaluation of methods to assign cell type labels to cell clusters from single-cell RNA-sequencing data. F1000Research. 2019 Oct 14;8:296. https://pubmed.ncbi.nlm.nih.gov/31508207/

50. Mikhailov N, Leskinen J, Fagerlund I, Poguzhelskaya E, Giniatullina R, Gafurov O, et al. Mechanosensitive meningeal nociception via Piezo channels: Implications for pulsatile pain in migraine? Neuropharmacology. 2019 May;149:113–23.

51. Velasco-Estevez M, Gadalla KKE, Liñan-Barba N, Cobb S, Dev KK, Sheridan GK. Inhibition of Piezo1 attenuates demyelination in the central nervous system. Glia. 2020 Feb;68(2):356–75.

52. Konttinen H, Cabral-da-Silva Me. Cmecme. C, Ohtonen S, Wojciechowski S, Shakirzyanova A, Caligola S, et al. PSEN1ΔE9, APPswe, and APOE4 confer disparate phenotypes in human iPSC-derived microglia. Stem Cell Reports. 2019 Oct 8;13(4):669–83.

53. Zhang Y, Sloan SA, Clarke LE, Caneda C, Plaza CA, Blumenthal PD, et al. Purification and Characterization of Progenitor and Mature Human Astrocytes Reveals Transcriptional and Functional Differences with Mouse. Neuron. 2016 Jan 6;89(1):37–53.

54. Zhang Y, Chen K, Sloan SA, Bennett ML, Scholze AR, O’Keeffe S, et al. An RNA-sequencing transcriptome and splicing database of glia, neurons, and vascular cells of the cerebral cortex. J Neurosci. 2014 Sep 3;34(36):11929–47.

55. Gosselin D, Skola D, Coufal NG, Holtman IR, Schlachetzki JCM, Sajti E, et al. An environment-dependent transcriptional network specifies human microglia identity. Science (80-). 2017 Jun 23;356(6344):1248–59.

56. Srinivasan K, Friedman BA, Etxeberria A, Huntley MA, van der Brug MP, Foreman O, et al. Alzheimer’s Patient Microglia Exhibit Enhanced Aging and Unique Transcriptional Activation. Cell Rep. 2020 Jun 30;31(13):107843.

57. Mathys H, Davila-Velderrain J, Peng Z, Gao F, Mohammadi S, Young JZ, et al. Single-cell transcriptomic analysis of Alzheimer’s disease. Nature. 2019 Jun 1;570(7761):332–7. http://www.nature.com/articles/s41586-019-1195-2

58. Galatro TF, Holtman IR, Lerario AM, Vainchtein ID, Brouwer N, Sola PR, et al. Transcriptomic analysis of purified human cortical microglia reveals age-associated changes. Nat Neurosci. 2017 Aug 1;20(8):1162–71.

59. Syeda R, Xu J, Dubin AE, Coste B, Mathur J, Huynh T, et al. Chemical activation of the mechanotransduction channel Piezo1. Elife. 2015;4(MAY).

60. Bae C, Sachs F, Gottlieb PA. The mechanosensitive ion channel Piezo1 is inhibited by the peptide GsMTx4. Biochemistry. 2011 Jul 26;50(29):6295–300. https://pubmed.ncbi.nlm.nih.gov/21696149/

61. Montilla A, Zabala A, Matute C, Domercq M. Functional and Metabolic Characterization of Microglia Culture in a Defined Medium. Front Cell Neurosci. 2020 Feb 7;14:22. https://www.frontiersin.org/article/10.3389/fncel.2020.00022/full

62. Ayata P, Schaefer A. Innate sensing of mechanical properties of brain tissue by microglia. Vol. 62, Current Opinion in Immunology. Elsevier Ltd; 2020. p. 123–30.

63. Nunes P, Demaurex N. The role of calcium signaling in phagocytosis. J Leukoc Biol. 2010 Jul;88(1):57–68.

64. Podleśny-Drabiniok A, Marcora E, Goate AM. Microglial Phagocytosis: A Disease-Associated Process Emerging from Alzheimer’s Disease Genetics. Vol. 43, Trends in Neurosciences. Elsevier Ltd; 2020. p. 965–79.

65. Novikova G, Kapoor M, Tcw J, Abud EM, Efthymiou AG, Chen SX, et al. Integration of Alzheimer’s disease genetics and myeloid genomics identifies disease risk regulatory elements and genes. Nat Commun. 2021 Dec 1;12(1):1–14. https://doi.org/10.1038/s41467-021-21823-y

66. Sheng J, Su L, Xu Z, Chen G. Progranulin polymorphism rs5848 is associated with increased risk of Alzheimer’s disease. Gene. 2014 Jun;542(2):141–5.

67. Brawek B, Schwendele B, Riester K, Kohsaka S, Lerdkrai C, Liang Y, et al. Impairment of in vivo calcium signaling in amyloid plaque-associated microglia. Acta Neuropathol. 2014 Jan;127(4):495–505.

68. Savchenko E, Malm T, Konttinen H, Hämäläinen RH, Guerrero-Toro C, Wojciechowski S, et al. Aβ and inflammatory stimulus activate diverse signaling pathways in monocytic cells: Implications in retaining phagocytosis in Aβ-laden environment. Front Cell Neurosci. 2016 Dec;10(DEC2016).

69. McLarnon JG, Choi HB, Lue LF, Walker DG, Kim SU. Perturbations in calcium-mediated signal transduction in microglia from Alzheimer’s disease patients. J Neurosci Res. 2005 Aug;81(3):426– 35.

70. Esparza TJ, Wildburger NC, Jiang H, Gangolli M, Cairns NJ, Bateman RJ, et al. Soluble amyloid-beta aggregates from human Alzheimer’s disease brains. Sci Rep. 2016 Dec;6.

71. Malmberg M, Malm T, Gustafsson O, Sturchio A, Graff C, Espay AJ, et al. Disentangling the Amyloid Pathways: A Mechanistic Approach to Etiology. Front Neurosci. 2020 Apr;14.

72. Malm T, Ort M, Tähtivaara L, Jukarainen N, Goldsteins G, Puoliväli J, et al. β-amyloid infusion results in delayed and age-dependent learning deficits without role of inflammation or β-amyloid deposits. Proc Natl Acad Sci U S A. 2006 Jun;103(23):8852–7.

73. Chen GF, Xu TH, Yan Y, Zhou YR, Jiang Y, Melcher K, et al. Amyloid beta: Structure, biology and structure-based therapeutic development. Vol. 38, Acta Pharmacologica Sinica. Nature Publishing Group; 2017. p. 1205–35.

74. Tabaton M, Piccini A. Role of water-soluble amyloid-b in the pathogenesis of Alzheimer’s disease.

75. Richard BC, Kurdakova A, Baches S, Bayer TA, Weggen S, Wirths O. Gene dosage dependent aggravation of the neurological phenotype in the 5XFAD mouse model of Alzheimer’s disease. J Alzheimer’s Dis. 2015 Jan;45(4):1223–36.

76. Jawhar S, Trawicka A, Jenneckens C, Bayer TA, Wirths O. Motor deficits, neuron loss, and reduced anxiety coinciding with axonal degeneration and intraneuronal Aβ aggregation in the 5XFAD mouse model of Alzheimer’s disease. Neurobiol Aging. 2012;33(1):196.e29-196.e40.

77. Unger MS, Schernthaner P, Marschallinger J, Mrowetz H, Aigner L. Microglia prevent peripheral immune cell invasion and promote an anti-inflammatory environment in the brain of APP-PS1 transgenic mice. J Neuroinflammation. 2018 Sep;15(1).

78. Heneka MT, Carson MJ, Khoury J El, Landreth GE, Brosseron F, Feinstein DL, et al. Neuroinflammation in Alzheimer’s disease. Vol. 14, The Lancet Neurology. Lancet Publishing Group; 2015. p. 388– 405.

79. Krabbe G, Halle A, Matyash V, Rinnenthal JL, Eom GD, Bernhardt U, et al. Functional Impairment of Microglia Coincides with Beta-Amyloid Deposition in Mice with Alzheimer-Like Pathology. PLoS One. 2013 Apr;8(4):60921.

80. Perlmutter L, Scott S, Barron E, Chui H. MHC Class 11-Positive Microglia in Human Brain: Association With Alzheimer Lesions. Vol. 33549458, Journal of Neuroscience Research. 1992.

81. Galloway DA, Phillips AEM, Owen DRJ, Moore CS. Phagocytosis in the brain: Homeostasis and disease. Vol. 10, Frontiers in Immunology. Frontiers Media S.A.; 2019. p. 790.

82. Qiu Z, Guo J, Kala S, Zhu J, Xian Q, Qiu W, et al. The Mechanosensitive Ion Channel Piezo1 Significantly Mediates In Vitro Ultrasonic Stimulation of Neurons. iScience. 2019;21:448–57.

83. Bobola MS, Chen L, Ezeokeke CK, Olmstead TA, Nguyen C, Sahota A, et al. Transcranial focused ultrasound, pulsed at 40 Hz, activates microglia acutely and reduces Aβ load chronically, as demonstrated in vivo. Brain Stimul. 2020;13(4):1014–23.

84. Gottlieb PA, Sachs F. Piezo1: Properties of a cation selective mechanical channel. Vol. 6, Channels. Taylor and Francis Inc.; 2012. p. 214.

85. Bae C, Gottlieb PA, Sachs F. Human PIEZO1: Removing inactivation. Biophys J. 2013 Aug;105(4):880–6.

86. Maroto R, Kurosky A, Hamill OP. Mechanosensitive Ca2+ permeant cation channels in human prostate tumor cells. Channels. 2012;6(4).

87. Hamill OP, Mcbride DW. Mechanogated channels in Xenopus oocytes: Different gating modes enable a channel to switch from a phasic to a tonic mechanotransducer. In: Biological Bulletin. Marine Biological Laboratory; 1997. p. 121–4.

88. Tsuchiya M, Hara Y, Okuda M, Itoh K, Nishioka R, Shiomi A, et al. Cell surface flip-flop of phosphatidylserine is critical for PIEZO1-mediated myotube formation. Nat Commun. 2018 Dec 1;9(1). https://pubmed.ncbi.nlm.nih.gov/29799007/

89. Bohlen CJ, Bennett FC, Tucker AF, Collins HY, Mulinyawe SB, Barres BA. Diverse Requirements for Microglial Survival, Specification, and Function Revealed by Defined-Medium Cultures. Neuron. 2017 May;94(4):759-773.e8. https://linkinghub.elsevier.com/retrieve/pii/S0896627317304002

90. Ridone P, Pandzic E, Vassalli M, Cox CD, Macmillan A, Gottlieb PA, et al. Disruption of membrane cholesterol organization impairs the activity of PIEZO1 channel clusters. J Gen Physiol. 2020 Aug 3;152(8). https://doi.org/10.1085/jgp.201912515

91. Xu Y ping, Zhang J wen, Li L, Ye Z you, Zhang Y, Gao X, et al. Complex regulation of capsaicin on intracellular second messengers by calcium dependent and independent mechanisms in primary sensory neurons. Neurosci Lett. 2012 May;517(1):30–5.

92. Poole K, Herget R, Lapatsina L, Ngo HD, Lewin GR. Tuning Piezo ion channels to detect molecular-scale movements relevant for fine touch. Nat Commun. 2014 Mar;5.

93. Qi Y, Andolfi L, Frattini F, Mayer F, Lazzarino M, Hu J. Membrane stiffening by STOML3 facilitates mechanosensation in sensory neurons. Nat Commun. 2015 Oct;6.

94. Tcw J, Liang S, Qian L, Pipalia N, Chao M, Shi Y, et al. Cholesterol and Matrisome Pathways Dysregulated in Human APOE ∈4 Glia. SSRN Electron J. 2019;

95. Lukacs V, Mathur J, Mao R, Bayrak-Toydemir P, Procter M, Cahalan SM, et al. Impaired PIEZO1 function in patients with a novel autosomal recessive congenital lymphatic dysplasia. Nat Commun. 2015 Sep;6.

96. Ma S, Cahalan S, LaMonte G, Grubaugh ND, Zeng W, Murthy SE, et al. Common PIEZO1 Allele in African Populations Causes RBC Dehydration and Attenuates Plasmodium Infection. Cell. 2018 Apr;173(2):443-455.e12.

97. Nguetse CN, Purington N, Ebel ER, Shakya B, Tetard M, Kremsner PG, et al. A common polymorphism in the mechanosensitive ion channel PIEZO1 is associated with protection from severe malaria in humans. Proc Natl Acad Sci U S A. 2020 Apr;117(16):9074–81.

98. Ma S, Dubin AE, Zhang Y, Mousavi SAR, Wang Y, Coombs AM, et al. A role of PIEZO1 in iron metabolism in mice and humans. Cell. 2021 Feb 18;184(4):969-982.e13.

99. Pathak MM, Nourse JL, Tran T, Hwe J, Arulmoli J, Le DTT, et al. Stretch-activated ion channel Piezo1 directs lineage choice in human neural stem cells. Proc Natl Acad Sci U S A. 2014;111(45):16148–53.

100. Blumenthal NR, Hermanson O, Heimrich B, Shastri VP. Stochastic nanoroughness modulates neuron-astrocyte interactions and function via mechanosensing cation channels. Proc Natl Acad Sci U S A. 2014;111(45):16124–9.

101. Satoh K, Hata M, Takahara S, Tsuzaki H, Yokota H, Akatsu H, et al. A novel membrane protein, encoded by the gene covering KIAA0233, is transcriptionally induced in senile plaque-associated astrocytes. Brain Res. 2006 Sep 7;1108(1):19–27. https://pubmed.ncbi.nlm.nih.gov/16854388/

102. Liddelow SA, Guttenplan KA, Clarke LE, Bennett FC, Bohlen CJ, Schirmer L, et al. Neurotoxic reactive astrocytes are induced by activated microglia. Nature. 2017 Jan 18;541(7638):481–7. http://www.nature.com/doifinder/10.1038/nature21029

103. McCaughey T, Sanfilippo PG, Gooden GEC, Budden DM, Fan L, Fenwick E, et al. A global social media survey of attitudes to human genome editing. Vol. 18, Cell Stem Cell. Cell Press; 2016. p. 569–72.

104. Brand A, Allen L, Altman M, Hlava M, Scott J. Beyond authorship: Attribution, contribution, collaboration, and credit. Learn Publ. 2015 Apr 1;28(2):151–5. www.pnas.org/site/misc/iforc.pdf

