## Supplementary figures and tables for "Microglial amyloid beta clearance is driven by PIEZO1 channels"

**Additional file 1: Supplementary figures and tables.**

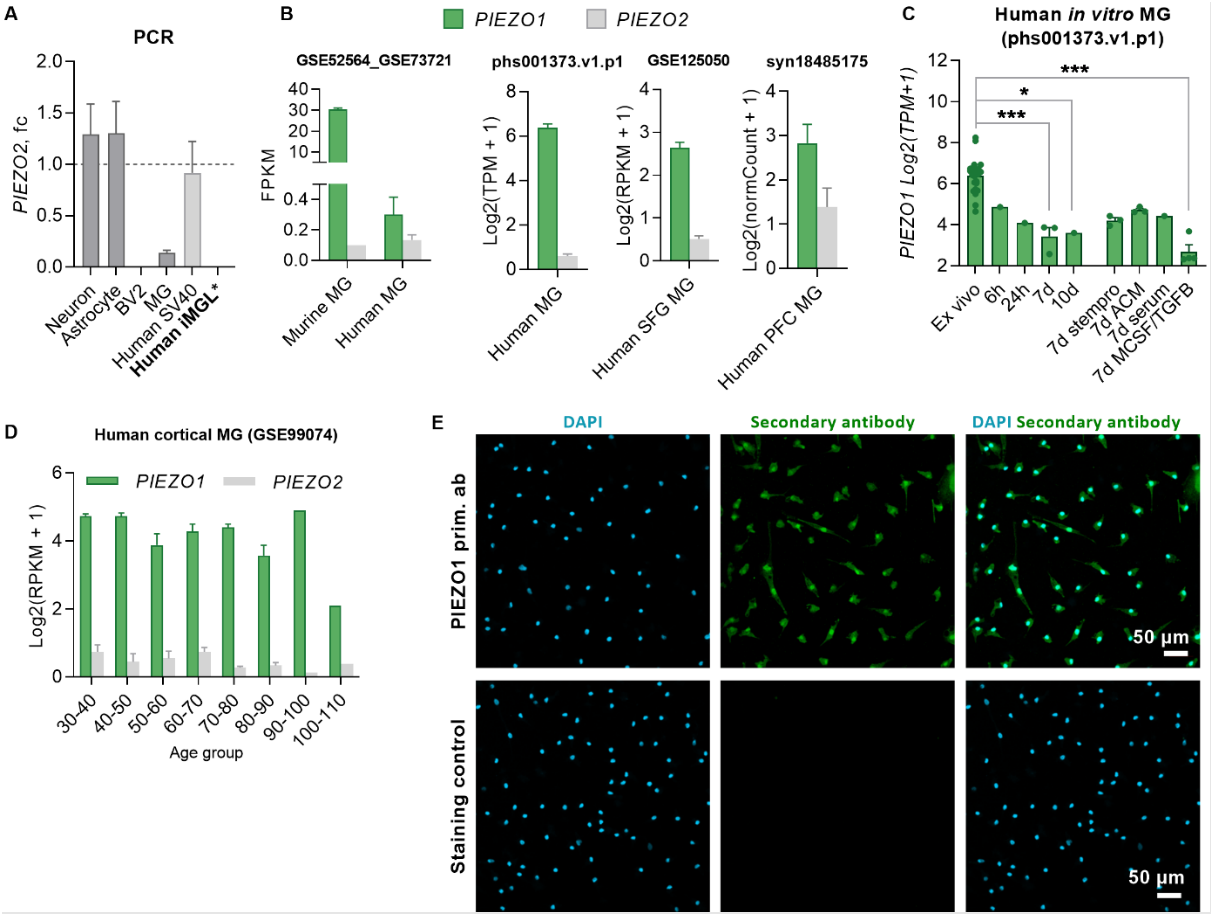


**Fig. S1. *PIEZO1* and *PIEZO2* expression in human and mouse RNA-seq datasets and staining controls for hiMGL immunostaining. A** *Piezo2* gene expression in murine trigeminal neurons (Neuron), astrocytes (Astro), microglia (MG) and microglial cell line (BV2); and in human microglial cell line (SV40) and iPSC-derived microglia (iMGL) analyzed by RT-qPCR (N=3-4). **B** *PIEZO1* and *PIEZO2* gene expression in microglia (MG) isolated from mouse (GSE52564) and human (GSE73721) brain, human neurosurgical brain tissue (phs001373.v1.p1), human postmortem superior frontal gyrus (SFG; GSE125050), and prefrontal cortex (PFC; syn18485175). **C** *PIEZO1* gene expression in RNA-seq data (phs001373.v1.p1) for *ex vivo* human microglia and cells cultured *in vitro* for up to ten days in medium supplemented with Stempro supplements, astrocyte conditioned medium (ACM) or with human serum, or MCSF and TGFB that are commonly used *in vitro* to support microglial identity. **D** *PIEZO1* and *PIEZO2* levels in superior parietal cortex at different ages (GSE99074). Data obtained for a-e from http://www.brainrnaseq.org/. **E** Immunostaining controls for DAPI (nuclei, blue) and PIEZO1 (green) antibody of *in vitro* iMGLs. Unpaired t-test ***p < 0.001, **p < 0.01, *p < 0.05; All data repeated in n experiments each with 3 replicates. Data as mean ± SEM. Exp=experiment; FPKM, Fragments Per Kilobase of transcript per Million; TPM, Transcripts Per Kilobase Million; RPKM Reads Per Kilobase Million.

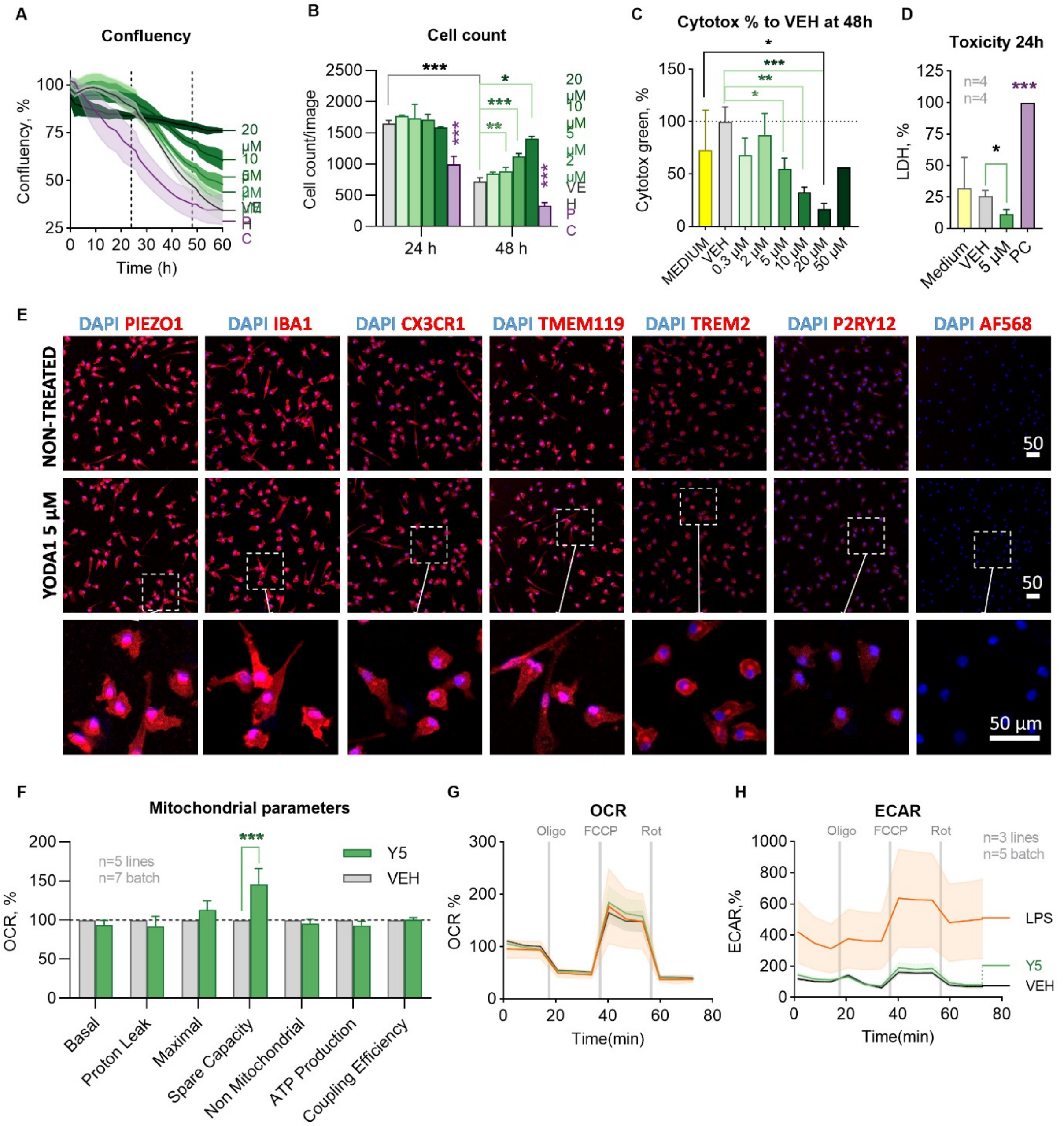


**Fig. S2. Activation of PIEZO1 orchestrates immune response of human iMGLs**. **A** Confluency of iMGLs in cytotox green assay over time (n=3). **B** Cell count at 24 h and 48 h (n=2 wells). **C** Quantification of cytotox green per confluence at 48 h show no differences for medium control, vehicle, 0.3 µM Yoda1 (Y0.3), 2 µM (Y2) Yoda1 (n=2) nor for 50 µM Yoda (n=1). Normalized to vehicle. **D** Lactate dehydrogenase (LDH) toxicity assay from iMGL cell culture medium at 24 h. N=2 for medium, others N=4 in n=4. **E** Representative images of human iMGLs immunostained after 24 h treatment with 5 μM Yoda1 labelled for antibodies for PIEZO1 and microglial markers IBA1 CX3CR1, TMEM119, TREM2 and P2RY12. As a staining control only-secondary-antibody AF568 was used. **F** All mitochondrial parameters calculated from OCR values of mitostress assay in fig 1 and normalized to vehicle. N=5 in n=7. **G** Oxygen consumption rate (OCR) of iMGLs in mitostress test with 20 ng/ml LPS, vehicle and 5 μM Yoda1. N=3 in n=5. **H** Corresponding extracellular asidification (ECAR) curve. All data repeated in n=experiments with N biological replicates and n≥3 technical replicates in each experiment. Unpaired t-test, one-way ANOVA or two-way ANOVA. Significance ***p < 0.001, **p < 0.01, *p < 0.05. Data as mean ± SEM

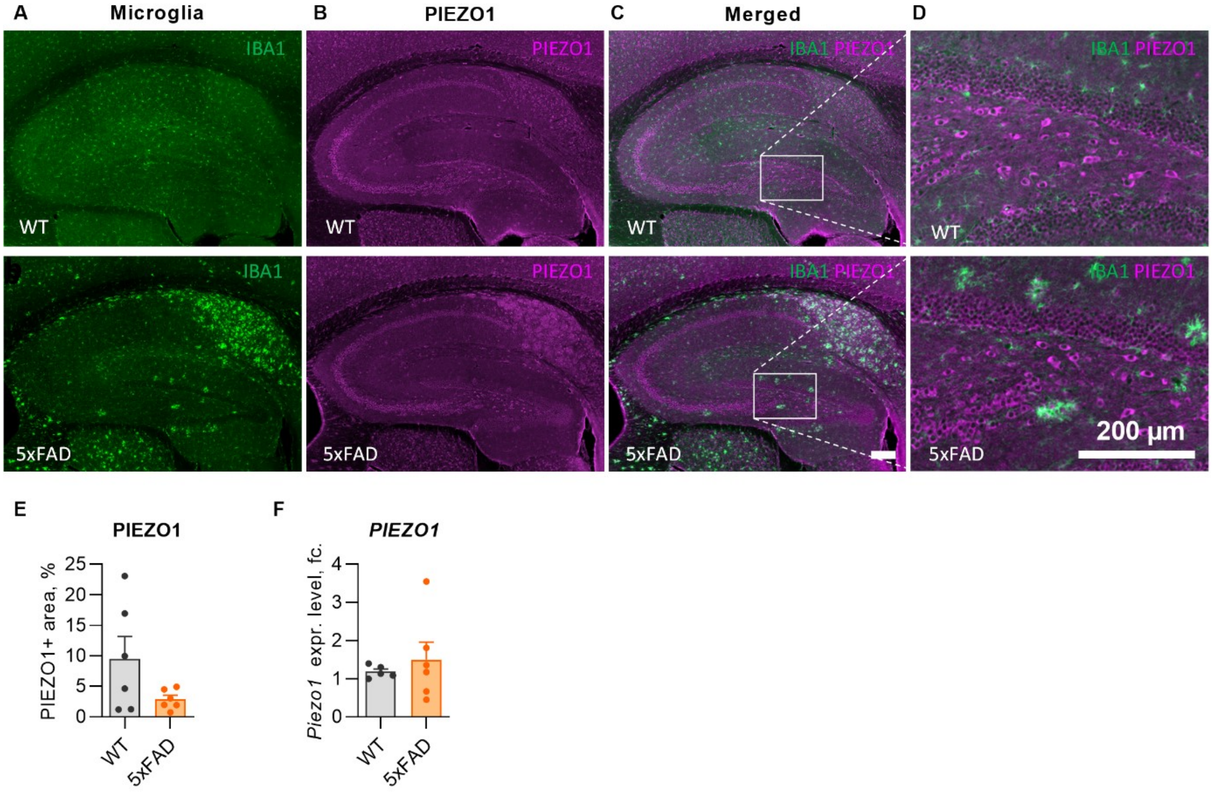


**Fig. S3. No differences in PIEZO1 expression in bulk brain tissue between WT and 5xFAD mice.** Representative immunofluorescence images of **A** microglia (IBA1, green), **B** PIEZO1 (magenta) and **C-D** merged channels from sagittal hippocampal sections of 5-month old WT and 5xFAD mice. Quantification of **E** immunoreactive area fraction in hippocampus and **F** gene expression in a hemisphere. N=6 mice. Data as mean ± SEM.

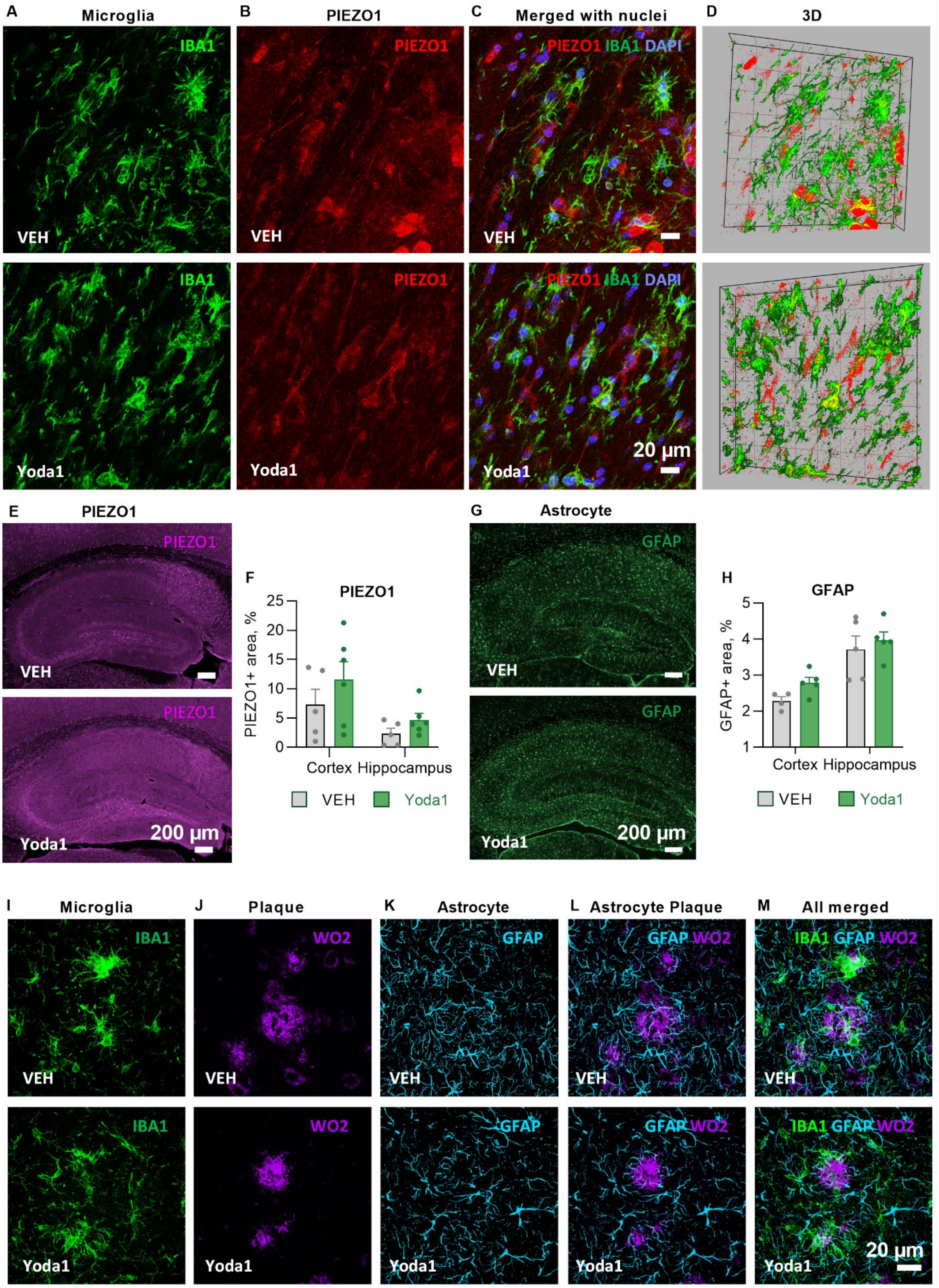


**Fig. S4. PIEZO1, microglia, astrocyte and Aβ plaque stainings in 5xFAD hippocampi.**

Representative images of maximum intensity projections of confocal z-stacks with staining's for **A** microglia (IBA1, green) and **B** PIEZO1 (red) with **C** merged channels showing co-localization of microglia and PIEZO1 (yellow) outside Aβ plaques in 5xFAD hippocampi. Scale bars 20 μm. **D** Corresponding 3D reconstructions of the z-stacks. **E** Representative immunofluorescence images of PIEZO1 (magenta) with **F** quantifications of immunoreactive area in hippocampus and cortex. Scale bar 200 μm. **G** Representative immunofluorescence images of astrocytes (GFAP, green) with **H** quantifications of immunoreactive area in hippocampus and cortex. Representative maximum intensity projections of confocal z-stack images of triple-immunostaining for **I** microglia (IBA1, green), **J** Aβ plaques (WO2, magenta), and **K** astrocytes (GFAP, green). **L** merged images showing no colocalization of GFAP and WO2 but **M** demonstrating clustering of IBA1 microglia around WO2 plaques. Scale bar 20 μm. N=5 VEH, N=6 Yoda1 mice. Unpaired t-test.

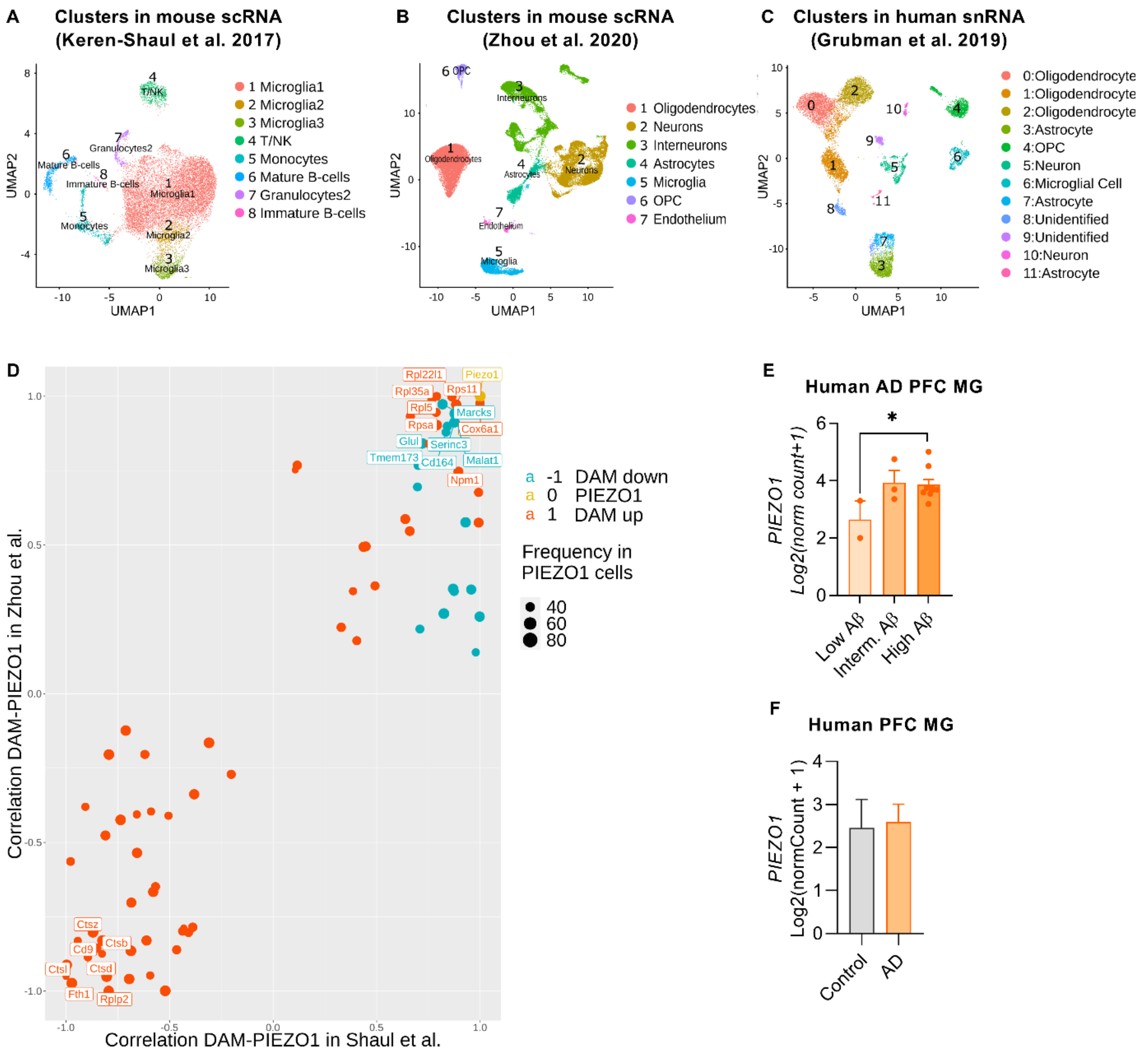


**Fig. S5. *PIEZO1* gene expression in published AD-related RNA datasets by our analysis.** UMAP visualizations of all cell clusters in **A** a 5xFAD mouse scRNA dataset [1], **B** Trem2$^{-/-}$ 5xFAD snRNA dataset [2], and **C** human AD patient entorhinal snRNA dataset [3] as result of our clustering and annotation. **D** A correlation diagram between *Piezo1* and the DAM signature genes in microglial subpopulations in mouse datasets [1,2]. Size of dot represents gene frequency in *Piezo1*+ cells, color down or upregulated in DAM microglia, and location depends by the correlation of the gene with *Piezo1*. *PIEZO1* expression in human microglia isolated from postmortem prefrontal cortex (PFC; syn18485175) of AD patients **E** with different levels of Aβ burden and **F** bulk data compared to healthy controls. The samples with zero *PIEZO1* expression were excluded. Data obtained for E-F from http://www.brainrnaseq.org/. Unpaired t-test. Significance *$p < 0.05$. Data as mean ± SEM or min and max. **See also Tables S1-3.**

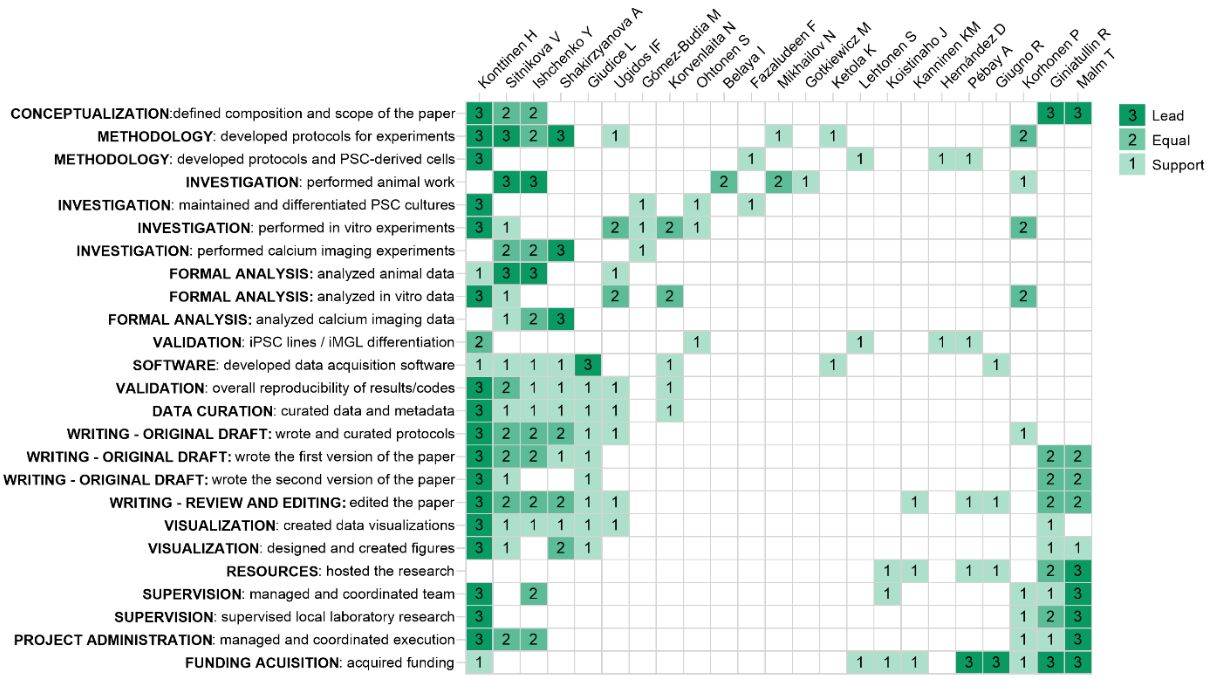


**Fig. S6. A diagram visualizing author contribution.** Based on the CRediT taxonomy [4]. For each type of contribution there are three levels indicated by color in the diagram: 1 support (light), 2 equal (medium), and 3 lead (dark).

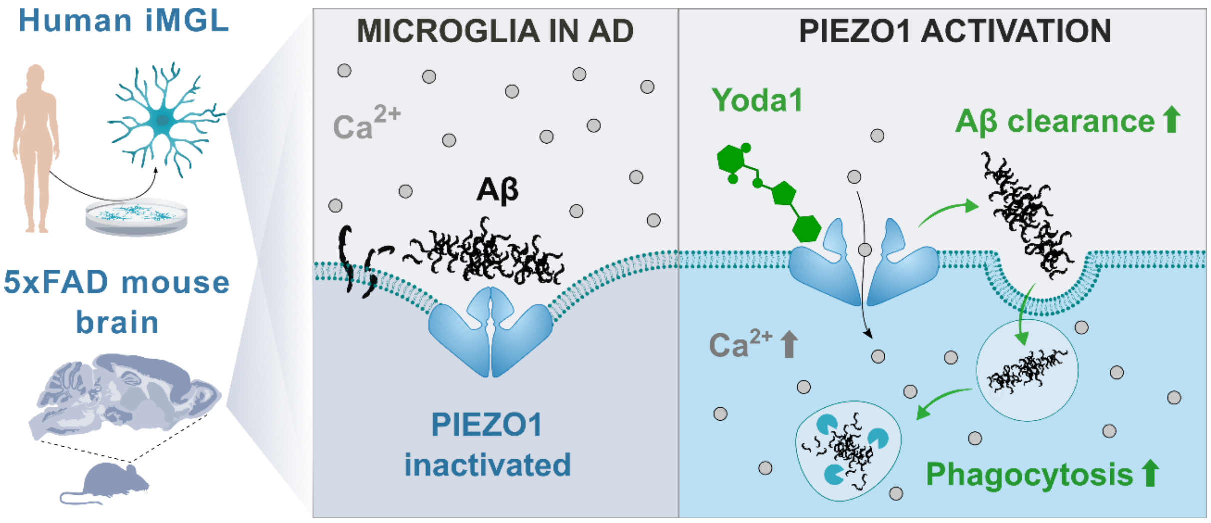


**Fig. S7. A graphical abstract summarizing the main finding of the paper.** Our data suggest that activating PIEZO1 mediated $Ca^{2+}$ influx with a selective agonist Yoda1 triggers microglia to shift their function in such manner that leads to phagocytosis and lysosomal activation in human iMGL and murine microglia *in vitro* which could explain the observed clearance of Aβ *in vivo* in response to administration of Yoda1 to 5xFAD mice.

**Table S1.** A correlation data for *Piezo1* and the DAM signature genes in microglial subpopulations in Keren-Shaul et al. 2017 dataset (GSE98969 [1]).

| Gene ID | Cor.Shaul | Microglia1 | Microglia2 | Microglia3 (DAM) | DAMs labels | min_freq.Shaul |
|---|---|---|---|---|---|---|
| ***Cd164*** | 0,84135765 | 0,992 | 0,974 | 0,965 | -1 | 0,965 |
| ***Cox6a1*** | 0,999093119 | 0,936 | 0,953 | 0,939 | 1 | 0,936 |
| ***Glul*** | 0,719356414 | 0,995 | 0,996 | 0,994 | -1 | 0,994 |
| ***Malat1*** | 0,87709347 | 0,997 | 0,996 | 0,965 | -1 | 0,965 |
| ***Marcks*** | 0,817983361 | 0,998 | 0,997 | 0,994 | -1 | 0,994 |
| ***Npm1*** | 0,894710871 | 0,949 | 0,897 | 0,985 | 1 | 0,897 |
| ***Piezo1*** | 1 | 1 | 0,998 | 0,997 | 0 | 0,997 |
| ***Rpl22l1*** | 0,862927676 | 0,835 | 0,725 | 0,939 | 1 | 0,725 |
| ***Rpl35a*** | 0,766646862 | 0,981 | 0,725 | 0,99 | 1 | 0,725 |
| ***Rpl5*** | 0,786998656 | 0,835 | 0,725 | 0,939 | 1 | 0,725 |
| ***Rps11*** | 0,883783624 | 0,98 | 0,984 | 0,985 | 1 | 0,98 |
| ***Rpsa*** | 0,790377515 | 0,975 | 0,984 | 0,977 | 1 | 0,975 |
| ***Serinc3*** | 0,875554683 | 0,999 | 0,998 | 0,994 | -1 | 0,994 |
| ***Tmem173*** | 0,70100146 | 0,983 | 0,989 | 0,939 | -1 | 0,939 |
| ***Cd9*** | -0,851741643 | 0,992 | 0,992 | 0,994 | 1 | 0,992 |
| ***Ctsb*** | -0,823464996 | 0,997 | 0,997 | 0,997 | 1 | 0,997 |
| ***Ctsd*** | -0,803587744 | 0,999 | 0,998 | 0,997 | 1 | 0,997 |
| ***Ctsl*** | -0,996532227 | 0,996 | 0,997 | 0,997 | 1 | 0,996 |
| ***Ctsz*** | -0,869983087 | 0,994 | 0,974 | 0,997 | 1 | 0,974 |
| ***Fth1*** | -0,972078623 | 0,996 | 0,996 | 0,997 | 1 | 0,996 |
| ***Rplp2*** | -0,793904002 | 0,982 | 0,984 | 0,985 | 1 | 0,982 |

DAM, disease associated microglia.

**Table S2.** A correlation data for *Piezo1* and the DAM signature genes in microglial subpopulations in Zhou et al. 2021 dataset (GSE140511 [2]).

| Gene ID | Cor.Zhou | 0 | 2 | 1 (DAM) | DAMs_labels.Zhou | min_freq.Zhou |
|---|---|---|---|---|---|---|
| *Cd164* | 0,899358766 | 0,93563 | 0,90831 | 0,74772 | -1 | 0,74772 |
| *Cox6a1* | 0,97677366 | 0,87632 | 0,75998 | 0,86958 | 1 | 0,75998 |
| *Glul* | 0,841592869 | 0,98637 | 0,98273 | 0,95384 | -1 | 0,95384 |
| *Malat1* | 0,94186021 | 0,99995 | 0,99995 | 0,99995 | -1 | 0,99995 |
| *Marcks* | 0,97312609 | 0,99286 | 0,90831 | 0,9781 | -1 | 0,90831 |
| *Npm1* | 0,746656312 | 0,85851 | 0,85309 | 0,88715 | 1 | 0,85309 |
| *Piezo1* | 1 | 0,99995 | 0,99995 | 0,99995 | 0 | 0,99995 |
| *Rpl22l1* | 0,998392023 | 0,90536 | 0,75998 | 0,90364 | 1 | 0,75998 |
| *Rpl35a* | 0,990164613 | 0,82883 | 0,85309 | 0,90364 | 1 | 0,82883 |
| *Rpl5* | 0,946034325 | 0,79143 | 0,90831 | 0,84561 | 1 | 0,79143 |
| *Rps11* | 0,972373749 | 0,90536 | 0,9405 | 0,94114 | 1 | 0,90536 |
| *Rpsa* | 0,903238077 | 0,91614 | 0,98273 | 0,96333 | 1 | 0,91614 |
| *Serinc3* | 0,912671379 | 0,99286 | 0,9405 | 0,98715 | -1 | 0,9405 |
| *Tmem173* | 0,767940761 | 0,93956 | 0,75998 | 0,78813 | -1 | 0,75998 |
| *Cd9* | -0,855908668 | 0,98883 | 0,75998 | 0,99803 | 1 | 0,75998 |
| *Ctsb* | -0,829803352 | 0,99414 | 0,99641 | 0,99916 | 1 | 0,99414 |
| *Ctsd* | -0,950962476 | 0,99646 | 0,9875 | 0,99956 | 1 | 0,9875 |
| *Ctsl* | -0,911484438 | 0,9719 | 0,9405 | 0,99183 | 1 | 0,9405 |
| *Ctsz* | -0,801830694 | 0,9809 | 0,98273 | 0,99724 | 1 | 0,9809 |
| *Fth1* | -0,973093154 | 0,9843 | 0,99729 | 0,98878 | 1 | 0,9843 |
| *Rplp2* | -0,998825952 | 0,89256 | 0,96043 | 0,90364 | 1 | 0,89256 |

DAM, disease associated microglia.

**Table S3.** Differentially expressed genes (DEGs) specific for m1-subcluster in snRNA Grubman et al. 2019 dataset (GSE138852, [3]). # indicates the eight DEGs expressed at highest level in m1 and *AD GWAS genes presented by Grubman et al. 2019.

| Gene ID | p_val | avg_logFC | pct.1 | pct.2 | p_val_adj | # | * |
|---|---|---|---|---|---|---|---|
| ***LINGO1*** | 6.12E-54 | 2.105339101 | 0.879 | 0.374 | 5.83971E-50 | # | |
| ***MT-ND4*** | 8.14E-18 | 1.745256134 | 0.561 | 0.15 | 7.77697E-14 | # | |
| ***MT-ND3*** | 5.7E-12 | 1.690972913 | 0.379 | 0.048 | 5.44469E-08 | | |
| ***BOK*** | 1.3E-10 | 1.597021176 | 0.242 | 0.012 | 1.2433E-06 | | |
| ***FCGBP*** | 7.95E-10 | 1.583308972 | 0.318 | 0.051 | 7.59101E-06 | | |
| ***CRYAB*** | 6.71E-11 | 1.567980192 | 0.394 | 0.147 | 6.41161E-07 | | |
| ***SNX6*** | 1.44E-09 | 1.497222753 | 0.348 | 0.09 | 1.37798E-05 | | |
| ***RPS28*** | 1.69E-13 | 1.494904224 | 0.485 | 0.135 | 1.61514E-09 | | |
| ***HSPA5*** | 4E-08 | 1.490987216 | 0.258 | 0.033 | 0.000381564 | | |
| ***MT-CO2*** | 1.99E-11 | 1.475309993 | 0.47 | 0.138 | 1.90464E-07 | # | |
| ***GFAP*** | 2.07E-08 | 1.433313829 | 0.288 | 0.051 | 0.000198083 | | |
| ***MT-ND2*** | 3.78E-08 | 1.432516142 | 0.333 | 0.078 | 0.000361407 | # | |
| ***NDRG1*** | 2.05E-07 | 1.400422926 | 0.212 | 0.021 | 0.001960356 | | |
| ***HSP90B1*** | 3.48E-07 | 1.394687521 | 0.318 | 0.078 | 0.003325934 | | |
| ***SPP1*** | 5.48E-12 | 1.349364779 | 0.773 | 0.455 | 5.23015E-08 | | |
| ***MT-ATP6*** | 2.97E-08 | 1.338138732 | 0.318 | 0.075 | 0.000283221 | # | |
| ***MT-CO3*** | 1.18E-11 | 1.311707022 | 0.53 | 0.129 | 1.12864E-07 | # | |
| ***MT-CYB*** | 1.59E-08 | 1.263223432 | 0.394 | 0.105 | 0.000152088 | # | |
| ***PLEKHA6*** | 4.56E-06 | 1.235822595 | 0.212 | 0.042 | 0.043528261 | | |
| ***DNAJB2*** | 3.31E-06 | 1.230809051 | 0.242 | 0.054 | 0.031584378 | | |
| ***LINC00486*** | 1.22E-39 | 1.204267915 | 1 | 0.979 | 1.16508E-35 | | |

| | | | | | | |
|---|---|---|---|---|---|---|
| ***APOC1*** | 1.31E-06 | 1.177523416 | 0.288 | 0.099 | 0.01249037 | * |
| ***RPL35*** | 6.29E-07 | 1.102400971 | 0.394 | 0.144 | 0.006008747 | |
| ***VWA1*** | 3.94E-06 | 1.101675686 | 0.152 | 0.009 | 0.037653863 | |
| ***ARMC9*** | 5.39E-09 | 1.088684428 | 0.515 | 0.231 | 5.14294E-05 | |
| ***HSPA1A*** | 1.16E-08 | 1.071470619 | 0.545 | 0.189 | 0.000110632 | |
| ***KANSL1L*** | 4.29E-06 | 1.067163753 | 0.258 | 0.078 | 0.040951065 | |
| ***DPYD*** | 6.98E-07 | 1.03478417 | 0.5 | 0.266 | 0.006662666 | |
| ***HSP90AA1*** | 3.51E-07 | 1.032117928 | 0.439 | 0.141 | 0.003354925 | |
| ***RPL37A*** | 2.23E-06 | 1.000686264 | 0.303 | 0.111 | 0.021285833 | |
| ***RPS19*** | 2.54E-07 | 0.971602862 | 0.53 | 0.341 | 0.002423795 | |
| ***C1QC*** | 6.19E-08 | 0.899225936 | 0.455 | 0.263 | 0.000591165 | |
| ***FMN1*** | 9.03E-08 | 0.898793765 | 0.5 | 0.251 | 0.000862091 | |
| ***ACTB*** | 6.81E-07 | 0.870135708 | 0.53 | 0.311 | 0.006503316 | |
| ***SERF2*** | 1.72E-07 | 0.869088241 | 0.333 | 0.186 | 0.001645651 | |
| ***APOE*** | 8.72E-08 | 0.822610932 | 0.697 | 0.533 | 0.000832737 | * |
| ***RFWD2*** | 4.32E-06 | 0.329535712 | 0.227 | 0.249 | 0.041250532 | |
| ***UBE2E2*** | 1.49E-07 | -0.265467932 | 0.364 | 0.683 | 0.001421866 | |
| ***KCNMA1*** | 2.59E-07 | -0.37860586 | 0.167 | 0.428 | 0.002471759 | |
| ***AOAH*** | 2.31E-07 | -0.458036774 | 0.197 | 0.494 | 0.00220782 | |
| ***HS3ST4*** | 3.71E-06 | -0.490835876 | 0.379 | 0.701 | 0.035405419 | |
| ***MEF2A*** | 1.27E-08 | -0.537339438 | 0.545 | 0.874 | 0.000120936 | |
| ***TMEM117*** | 1.29E-06 | -0.53980123 | 0.015 | 0.156 | 0.012358666 | |
| ***ANKRD44*** | 2.35E-09 | -0.582183292 | 0.318 | 0.722 | 2.24312E-05 | |
| ***MEF2C*** | 7.89E-08 | -0.593242864 | 0.576 | 0.88 | 0.000753268 | * |
| ***C10orf11*** | 1.2E-14 | -0.704657839 | 0.606 | 0.958 | 1.14534E-10 | |
| ***MALAT1*** | 1.51E-39 | -0.719913013 | 1 | 1 | 1.44389E-35 | |
| ***FOXN3*** | 8.51E-09 | -0.727831377 | 0.364 | 0.757 | 8.12227E-05 | * |

| | | | | | | |
|---|---|---|---|---|---|---|
| ***DOCK4*** | 2.11E-09 | -0.757036602 | 0.758 | 0.949 | 2.01274E-05 | |
| ***ST6GAL1*** | 1.54E-09 | -0.876638131 | 0.439 | 0.817 | 1.46945E-05 | * |
| ***CAMK1D*** | 2.54E-06 | -1.093782153 | 0.061 | 0.335 | 0.024263186 | |
| ***LPAR6*** | 6.78E-09 | -1.109260972 | 0.227 | 0.629 | 6.47025E-05 | |
| ***USP53*** | 1.34E-06 | -1.119665536 | 0.045 | 0.311 | 0.012819522 | |
| ***USP39*** | 7.13E-07 | -1.161382456 | 0.076 | 0.38 | 0.00681296 | |
| ***ZFP36L2*** | 5.21E-07 | -1.171247383 | 0.045 | 0.323 | 0.004977081 | |
| ***A2M*** | 1.69E-06 | -1.173381626 | 0.182 | 0.497 | 0.016146244 | |
| ***SYNDIG1*** | 7.3E-09 | -1.259017784 | 0.167 | 0.56 | 6.96651E-05 | |
| ***CD86*** | 3.4E-06 | -1.305477311 | 0.076 | 0.356 | 0.032499009 | |
| ***BTG2*** | 2.53E-06 | -1.323070172 | 0 | 0.177 | 0.024127191 | |
| ***RP11-624C23.1*** | 5.72E-16 | -1.337719226 | 0.197 | 0.74 | 5.46382E-12 | |
| ***PRDM11*** | 2.67E-07 | -1.363609996 | 0 | 0.207 | 0.002552475 | |
| ***APBA1*** | 1.07E-07 | -1.379359477 | 0 | 0.219 | 0.001020117 | |
| ***FRMD4A*** | 1.03E-22 | -1.382930811 | 0.439 | 0.919 | 9.87037E-19 | * |
| ***OLR1*** | 9.56E-07 | -1.405655026 | 0.106 | 0.404 | 0.009130846 | |
| ***KCNQ1*** | 2.64E-08 | -1.470846884 | 0 | 0.237 | 0.000252285 | * |
| ***SDK1*** | 1.15E-08 | -1.471588901 | 0.106 | 0.467 | 0.000109763 | |
| ***CSGALNACT1*** | 4.54E-10 | -1.565128643 | 0.061 | 0.446 | 4.33123E-06 | |
| ***MCF2L2*** | 1.74E-07 | -1.576547828 | 0.045 | 0.344 | 0.001657942 | |
| ***ST6GALNAC3*** | 4.49E-13 | -1.578137257 | 0.106 | 0.581 | 4.2879E-09 | |
| ***RP5-1031D4.2*** | 1.07E-07 | -1.608265176 | 0 | 0.219 | 0.001020117 | |
| ***RP11-480C22.1*** | 2.09E-08 | -1.628153071 | 0 | 0.24 | 0.000199362 | |
| ***DUSP1*** | 1.07E-07 | -1.758573775 | 0 | 0.219 | 0.001020117 | |
| ***SRGN*** | 4.29E-10 | -2.034698722 | 0.076 | 0.458 | 4.09934E-06 | |

**References**


1. Keren-Shaul H, Spinrad A, Weiner A, Matcovitch-Natan O, Dvir-Szternfeld R, Ulland TK, et al. A Unique Microglia Type Associated with Restricting Development of Alzheimer's Disease. Cell. Cell Press; 2017;169:1276-1290.e17.

2. Zhou Y, Song WM, Andhey PS, Swain A, Levy T, Miller KR, et al. Human and mouse single-nucleus transcriptomics reveal TREM2-dependent and TREM2-independent cellular responses in Alzheimer's disease. Nat Med [Internet]. Nature Research; 2020 [cited 2021 May 20];26:131–42. Available from: http://www.nature.com/articles/s41591-019-0695-9

3. Grubman A, Chew G, Ouyang JF, Sun G, Choo XY, McLean C, et al. A single-cell atlas of entorhinal cortex from individuals with Alzheimer's disease reveals cell-type-specific gene expression regulation. Nat Neurosci. Nature Research; 2019;22:2087–97.

4. Brand A, Allen L, Altman M, Hlava M, Scott J. Beyond authorship: Attribution, contribution, collaboration, and credit. Learn Publ [Internet]. Association of Learned and Professional Society Publ.; 2015 [cited 2021 Jul 6];28:151–5. Available from: www.pnas.org/site/misc/iforc.pdf
